# Human and murine neutrophils share core transcriptional programs in both homeostatic and inflamed contexts

**DOI:** 10.1101/2022.11.13.516246

**Authors:** Nicolaj S. Hackert, Felix A. Radtke, Tarik Exner, Hanns-Martin Lorenz, Carsten Müller-Tidow, Peter A. Nigrovic, Guido Wabnitz, Ricardo Grieshaber-Bouyer

## Abstract

Neutrophils are frequently studied in murine models, but the extent to which findings translate to humans remains poorly defined. Here, we performed an integrative transcriptomic analysis of 11 murine and 13 human datasets. In homeostasis, neutrophils exhibited the highest number of lineage-specific genes and the greatest degree of correlated expression among genes with one-to-one orthologs (*r* = 0.79, *P* < 2.2 × 10^−16^) compared to other leukocytes. In inflammation, neutrophils displayed considerable transcriptional diversity, but shared a core inflammation program across a broad range of conditions which was conserved between species. This core program included genes encoding IL-1 family members, CD14, IL-4R, CD69 and PD-L1. Chromatin accessibility of core inflammation genes increased significantly in blood compared to bone marrow and further with migration from blood to tissue. Transcription factor enrichment analysis nominated members of the NF-κB family and AP-1 complex as important drivers of the core inflammation program, and HoxB8 neutrophils with JUNB knockout showed a significantly reduced expression of core inflammation genes at baseline and upon stimulation. In vitro perturbations confirmed surface protein upregulation of core inflammation members in both species. Together, we demonstrate substantial transcriptional conservation in neutrophils in homeostasis and identify a core inflammation program conserved across species. This systems biology approach can be leveraged to improve transitions between the murine and human context.

**Key Points:**

1. The transcriptome of resting neutrophils is substantially conserved between humans and mice
2. A core inflammation program in neutrophils is shared across a broad range of conditions and conserved across humans and mice

**Graphical Abstract:** 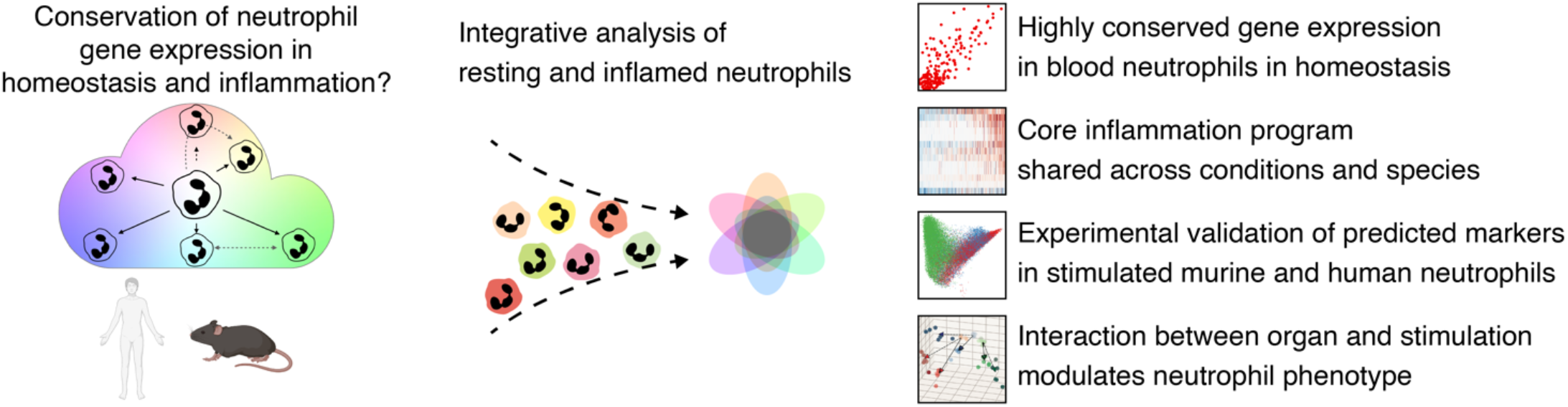

## INTRODUCTION

Neutrophils mediate homeostatic and inflammatory processes and display substantial phenotypic and functional heterogeneity. While animal models fuel fundamental discoveries in hematopoiesis, differences between humans and mice can impair the translation of findings (1). To maximize impact on human health, life sciences increasingly benefit from seamless transitions between the murine and human system. However, due to structural and functional differences in genomes, it is often unclear which aspects reflect conserved biology. Therefore, integrative analyses of cellular systems across species are important for the success of translational research.

Structurally, the murine and human genome are closely related. They harbor ∼16,000 protein-coding genes considered to be one-to-one orthologs with high confidence (2). However, structural orthology does not equal functional similarity since expression patterns of orthologous genes can deviate substantially across organs and development (3). In leukocytes, expression of most orthologous genes and lineage-specific genes in particular, is well conserved between humans and mice (4). Despite this overall similarity, different species can display substantial differences in ortholog expression between tissues (5). For example, human neutrophils are highly abundant in defensins, yet their murine orthologs are expressed in gut epithelial cells, not in neutrophils.

Furthermore, neutrophils display high phenotypic and functional heterogeneity as a function of organ, maturation and inflammatory condition (6-9), but whether a core inflammation program consisting of genes that become induced across a range of inflammatory conditions exists, is not known. It is thus unclear how similarities and differences between human and mouse transcriptomes should be interpreted, particularly in the context of different inflammatory conditions.

To address these gaps in knowledge, we performed an integrative analysis of resting and inflamed leukocytes from humans and mice and assessed the degree of conservation of gene expression. We found that human and murine transcriptomes could be analyzed together, and that lineage-specific gene expression was closely related between humans and mice. Correlation of gene expression was particularly strong in blood neutrophils. In homeostasis, we identified distinct clusters of orthologous genes marked by high and low concordance, as well as abundantly expressed genes without one-to-one orthologs.

We further studied how the neutrophil transcriptome changes in inflammation, using a wide range of studies covering *in vitro* and *in vivo* inflammation as well as resting conditions in human (10-21) and mouse (12, 22-31). While transcriptional responses to different activating stimuli were heterogenous, we identified a core inflammation program in neutrophils conserved across species and conditions. This core inflammation program represents a group of genes preferentially upregulated in inflamed contexts compared to other genes. We predicted upstream regulators and found increasing accessibility of core inflammation program members in ATAC-seq. *JunB* ^-/-^ HoxB8 cells displayed a lower upregulation of core inflammation genes when stimulated with zymosan compared to wild-type cells.

Finally, we validated members of the core inflammation program on the protein level using flow cytometry of stimulated human and mouse neutrophils. These data also exposed an interplay between tissue of origin and stimulation in driving the phenotype of the neutrophil inflammatory response.

Our approach illustrates that multiple datasets of mouse and human gene expression data can be effectively combined to identify patterns shared across conditions and conserved across species. This approach can be transferred to other cell types and organisms to facilitate studies comparing gene expression across species.

## RESULTS

### Integrative analysis of leukocyte gene expression across species

To assess gene expression similarities and differences between human and mouse immune cells, we obtained bulk RNA-seq data from six sorted leukocyte lineages as part of the Haemopedia atlas (12, 32) **(Figure 1A)**. This dataset consisted of a total of 76 samples of T cells, B cells, dendritic cells, monocytes, NK cells and neutrophils. We then integrated gene expression matrices by mapping protein-coding, one-to-one orthologous genes with high confidence according to ENSEMBL (33).

**Figure 1.**
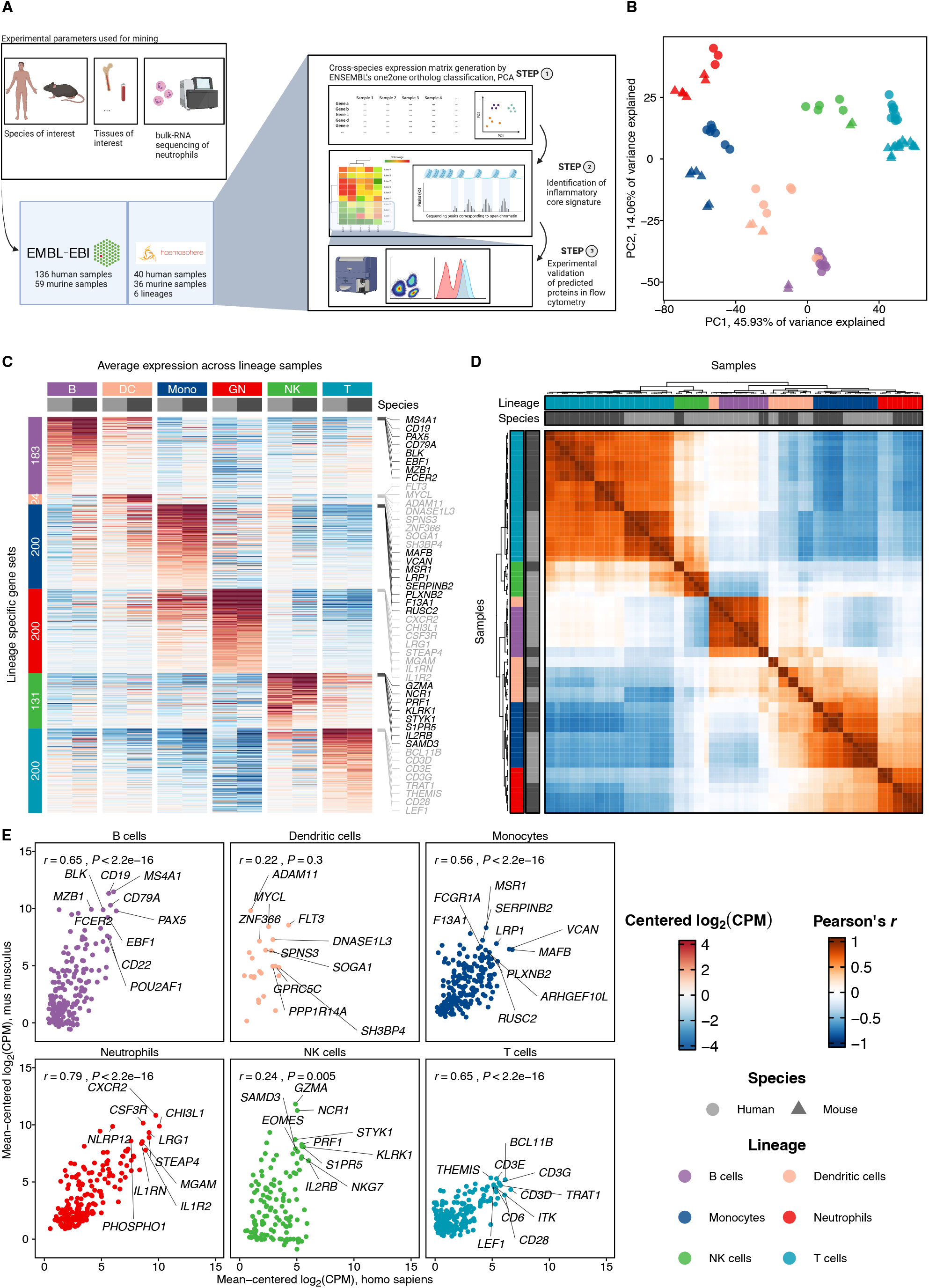
Integrative analysis of leukocyte gene expression across species. **(A)** Study overview. Gene expression from human and murine T cells, B cells, dendritic cells, monocytes, and neutrophils was obtained from the European Nucleotide Archive (ENA). Gene expression matrices were integrated using ENSEMBL high confidence orthologs of protein-coding genes. **(B)** Principal component analysis based on per-species mean-centered log_2_(CPM) of lineage-specific genes shows a distribution driven predominantly by lineage. **(C)** Concordant expression of the top lineage-specific genes for each lineage in each species. Shown is the average of log_2_(CPM) centered expression values in each lineage. Up to 200 genes are shown per lineage. For each lineage, 8 genes with the highest expression in the respective cell line are labeled. **(D)** Lineage-specific gene expression dominates the species effect. Shown is the clustering of Pearson’s *R* based on the centered expression of top lineage-specific genes. Neutrophils display the strongest correlation of lineage-specific gene expression across humans and mice compared to other leukocyte lineages. Gene expression (mean-centered log_2_(CPM)) of lineage-specific genes was defined as above. Correlation (Pearson’s *R*) between human (x) and mouse (y) gene expression is shown on the top left. The top 10 most abundantly expressed genes are labeled.

To evaluate the robustness of this approach, we performed a principal component analysis on the integrated expression matrix. For each lineage, up to 200 lineage-associated genes were selected as those with differential expression (log_2_ fold change > 0, *P*_adj_ ≤ 0.05) compared to all other lineages. Here, sample distribution was driven predominantly by lineage, followed by species **(Figure 1B)**. As envisioned, lineage-associated gene expression was highest in each respective lineage and occurred across species in all lineages **(Figure 1C)**. Similarly, clustering of sample-wise Pearson correlation coefficients based on these genes was driven predominantly by lineage, confirming that in our analytical approach, lineage identity dominates species differences **(Figure 1D)**.

Correspondingly, expression of key lineage-associated genes was highly conserved between humans and mice **(Figure 1E)**, for example *CSF3R* and *CHI3L1* in neutrophils, *CD19* and *CD22* in B cells, CD3 molecules and *CD28* in T cells, *NKG7* and *GZMA* in NK cells, *MSR1* and *SERPINB2* in monocytes and *FLT3* and *MYCL* in dendritic cells. The highest correlation between human and murine gene expression was observed in neutrophils (*r* = 0.79), followed by T-cells (0.65), B-cells (0.65), Monocytes (0.56), and a weaker correlation in NK cells (0.24) and dendritic cells (0.22) **(Figure 1E)**.

In conclusion, mapping one-to-one orthologs allows an integrated analysis of leukocyte transcriptomes across species to identify conserved and divergent expression patterns of structurally related genes.

### Transcriptional conservation in resting neutrophils

To systematically analyze which genes display similar and divergent expression across species, we integrated transcriptional profiles of resting (i.e. not activated) neutrophils available through the Sequence Read Archive (SRA). In a total of 84 human and 39 mouse samples, we observed high correlation in overall gene expression, transcription factor expression and lineage-associated gene expression across humans and mice (Pearson’s *r* between 0.78–0.87, *P* < 2.2 × 10^−16^) **(Figure 2A)**.

**Figure 2.**
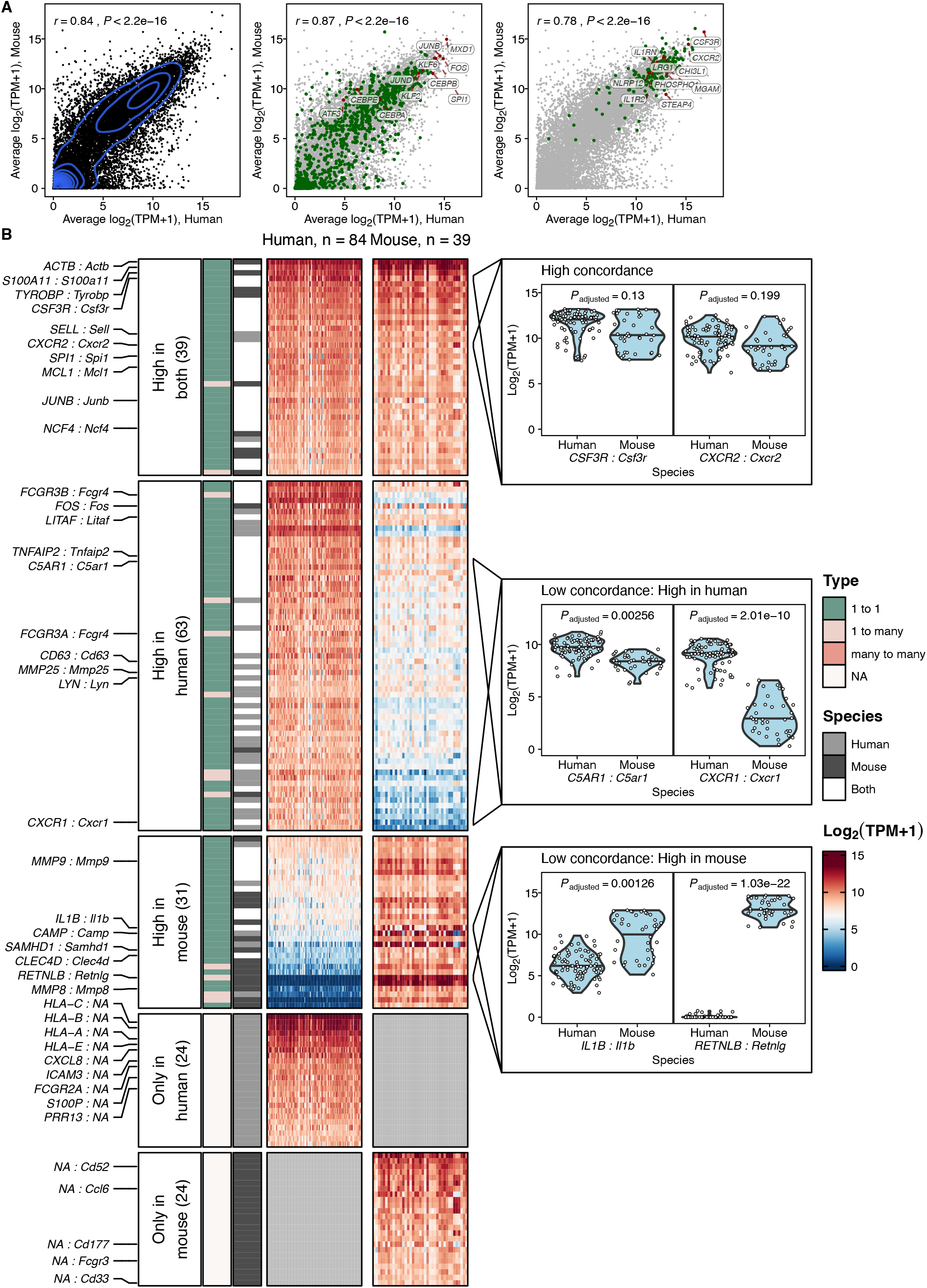
Conservation of neutrophil gene expression in homeostasis. Strong correlation of gene expression between resting human (x) and mouse (y) blood neutrophils. Left: all genes. Middle: transcription factors, highlighted in green. Transcription factors were retrieved from a curated set of transcription factors in ChEA3. The top 5 TFs (based on the sum of the average expression in human and mouse) were labeled and highlighted in red, additionally, we manually labeled and highlighted the genes *JUND, KLF2, ATF3, CEBPA, CEBPB* and *CEBPE*. Right: lineage-specific genes as depicted and defined in Figure 1, highlighted in green. Neutrophil genes were labeled as in Figure 1E and highlighted in red. Shown are log_2_(TPM+1) expression values. **(B)** Neutrophil lineage-associated genes with orthologs can show concordant or discordant expression across species. Gene expression heatmap (log_2_(TPM+1)) of neutrophil lineage-associated genes that were assigned to five different expression profile groups: high expression in both species, high expression in human/mouse and low in the other species, high in human/mouse and no high confidence ortholog; see Methods. Gene-gene pairs of particular importance in neutrophils are highlighted (HUMAN SYMBOL : Mouse symbol). Orthology relationships between the respective genes as well as species in which the gene was detected as lineage-associated are annotated. Right, violin plots of selected gene-gene-pairs showing their expression in individual samples for each species. Adjusted P-values from the linear mixed model used to define the groups are shown for each highlighted gene-gene pair.

We next focused on neutrophil lineage-associated genes and defined *GENE* : *Gene* (*HUMAN* : *Murine*) pairs that were assigned to one of five groups based on their expression patterns. For this step, we considered high confidence one-to-many and many-to-many orthologs in addition to one-to-one orthologs.

Orthologs with high expression in both humans and mice included the key neutrophil genes *CSF3R* (encoding the G-CSF receptor), *CXCR2, NCF4* (neutrophil cytosolic factor 4), and the transcription factors *MCL1, SPI1* (encoding PU.1, an essential transcription factor for terminal granulopoiesis (34, 35)) and *JUNB* **(Figure 2B)**. *JUNB*, a transcription factor prominently expressed in late neutrotime which plays a vital role in the inflammatory response of neutrophils (9, 36) **(Figure 2B)**. The concordance in expression of *CSF3R, CXCR2* and *JUNB* suggests that the analyzed neutrophils from humans and mice were of comparable developmental stage.

Orthologs with higher expression in human neutrophils included *FCGR3A* and *FCGR3B* (encoding CD16A and CD16B, respectively), which both are one-to-many orthologs of murine *Fcgr4*. This group also included the receptor for activated complement (*C5AR1*) and *CXCR1*, the receptor for CXCL8 (human)/KC (mouse). Genes with higher expression in mouse neutrophils included the protease *Mmp9, Camp* (encoding Cathelicidin Antimicrobial Peptide), *Il1b* and *Retnlg* (encoding Resistin-like gamma) **(Figure 2B)**.

Of note, most genes in categories 1–3 were one-to-one orthologs, although 13/133 (9.8 %) were one-to-many orthologs. Well-known neutrophil genes without one-to-one orthologs were also identified (categories 4 and 5) and included *CXCL8* in humans, a cytokine abundantly expressed in blood neutrophils and *Ccl6*, one of the most abundant chemokines in murine neutrophils **(Figure 2B)**. Enrichment for neutrophil-related GO terms was found across all five groups of genes **(Supplementary Figure S1)**.

In conclusion, human and murine neutrophils share substantial structural similarity in their expressed genes. Resting human and murine neutrophils display highly conserved expression of key neutrophil genes and transcription factors. Yet, several highly abundant neutrophil genes show divergent expression patterns or even lack structurally related counterparts in the other species, highlighting that both gene structure as well as species-specific expression need to be accounted in studies across species.

### A core inflammation program is shared across conditions and conserved across species

We next assessed how expression of one-to-one structural orthologs changes in different inflammatory contexts. Neutrophils display varied phenotypes in homeostasis and inflammation (6, 7, 9, 37), and a proportion of the transcriptional characteristics of different neutrophil states may be shared across different inflammatory conditions (9).

To identify changes in inflammation, we analyzed 11 studies encompassing a total of 46 resting and 66 activated neutrophil samples across different conditions **(Figure 3A, Supplementary Table 2)**. We tested for differential expression of genes with high-confidence one-to-one orthologs according to ENSEMBL separately within each study, comparing all reported conditions against their own resting controls and thereby isolating technical variation between studies.

**Figure 3.**
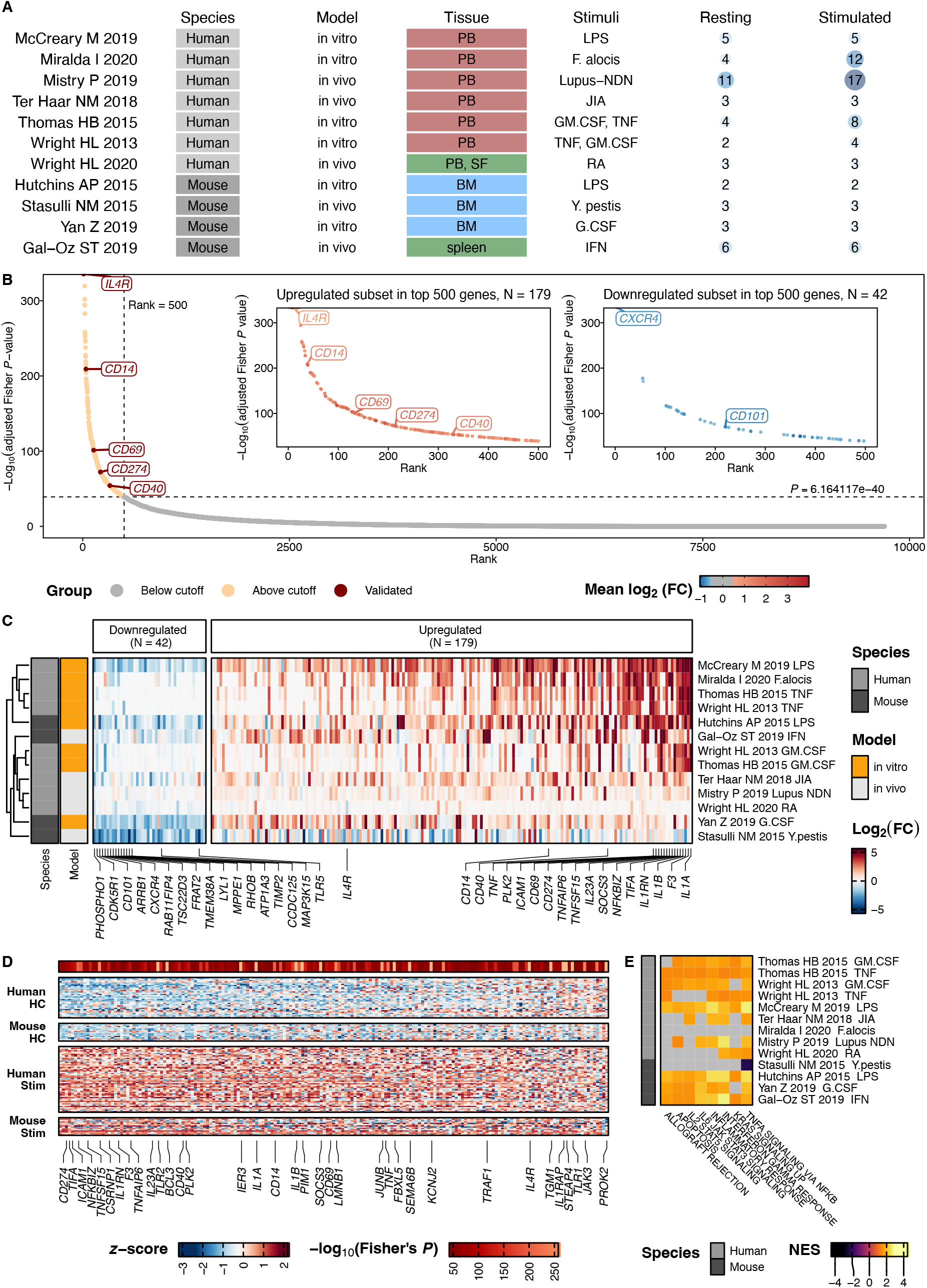
A core inflammation program is conserved across murine and human neutrophils. **(A)** Overview of the 11 studies integrated for analysis. Differential expression testing was performed independently for each study, so that resting neutrophils within each study were used as control. **(B)** A combined analysis of the neutrophil response to activation/inflammation identifies 179 consistently upregulated (core inflammation program) and 42 downregulated genes in inflammation. Shown are all (N = 9697) tested genes ranked by their −log_10_ adjusted Fisher *P*-value. The 500 genes with lowest *P*-value were subjected to an additional filtering step based on a log2 fold change cutoff ≥ 0.5 and ≤ −0.5 for upregulated and downregulated genes. Highlighted (*IL4R, CD14, CD69, CD274, CD40*) upregulated genes were validated experimentally (Figures 6–7). **(C)** 42 genes downregulated in inflammation and 179 core inflammation genes are shared across studies. Shown are the log_2_ fold changes across comparisons of genes up- and downregulated in inflammation. Rows represent a comparison, columns are genes that passed our meta-analysis thresholds. Columns are arranged by the mean log_2_ fold change across all comparisons. For each direction, the 15 genes with the highest absolute log_2_ fold change are labeled, as well as genes encoding for proteins validated in Figures 6–7. **(D)** Core inflammation genes are not expressed in resting neutrophils in both species and are induced upon activation. Shown is a heatmap with relative expression values (z-score for each gene across all samples) of the core inflammation genes. Each column represents a gene, each row a sample. *P*-values indicate the resultsy of an adjusted Fisher’s combined test. We labeled the top 20 genes with the lowest *P*-values, genes that were also labeled in Figure 3C, and manually labeled *TRAF1* and *JUNB*. **(E)** Conserved Gene Set Enrichment Analysis based on rankings derived from each comparison’s log_2_ fold changes. Heatmap showing normalized enrichment scores. Only pathways that have been significant in more than 50% of comparisons are depicted. Gray fields indicate non-significant NES values.

Compared to controls, inflamed neutrophils displayed 975 (median) differentially expressed genes (adjusted *P* < 0.05, absolute log_2_ fold change ≥ 0.5) **(Supplementary Figure 2A)**. These comprised 621 (median) significantly increased and 205 (median) significantly decreased genes. Both the number of differentially expressed genes and the genes themselves were highly heterogenous, indicating that neutrophils can undergo diverse transcriptional responses in inflammation.

We next searched for potential overlap in the inflammatory response shared across conditions. Such an overlap may represent a “core inflammation program”, from which neutrophils preferentially upregulate genes across a broad range of activating conditions. We used Fisher’s combined test to obtain a combined test statistic for each gene, summarizing individual comparisons from all datasets **(Supplementary Table 2)**. Based on the elbow of the *P*-value-rank plot, we selected from the top 500 genes with the lowest *P*-value those with absolute log_2_ fold change ≥ 0.5 **(Figure 3B)**.

A total of 221 genes displayed consistent changes in inflammation across studies: 179 genes were upregulated (the “core inflammation program”) and 42 genes were downregulated **(Figure 3C)**. Effect sizes of those 221 up- and downregulated genes agreed well across all tested comparisons and across species **(Figure 3C, Supplementary Figure 2B)**.

Core inflammation genes included the pro-inflammatory mediators *IL1A, IL1B, CD14, ICAM1, CD69, CD40, IL4R* and *CD274* (encoding PD-L1) **(Figure 3C)**.

Downregulated genes in inflammation included the cyclin-dependent kinase *CDK5R1, TLR5* (encoding Toll Like Receptor 5, an essential pathogen recognition receptor (38)), *CXCR4, CD101* and the member of the mitogen-activated protein kinase family *MAP3K15* **(Figure 3C)**.

As expression of CD101 and CXCR4 changes throughout neutrophil maturation and aging, we compared the fold change of these markers between neutrophils activated *in vitro* and those activated *in vivo*, to rule out effects of differential release from the bone marrow under stress. No differences were observed in either marker **(Supplementary Figure 2C)**, suggesting that the downregulation of CXCR4 and CD101 observed during neutrophil activation are cell-intrinsic and do not reflect a different maturation stage of neutrophils captured in the *in vivo* studies.

To further test the robustness of this core inflammation program, we performed an independent analysis using a linear mixed model. We observed a high replicability of our results, with differentially expressed genes (absolute β ≥ 1, *P*_adj_ < 0.05) identified by the linear mixed model showing a strong skewing towards low Fisher *P*-values and a π_1_-statistic of 0.55 **(Supplementary Figure 3)**.

We additionally assessed the replicability of differentially expressed genes between all tested comparisons. Median values of the π_1_-statistic ranged from 0.06 to 0.60, depending on the study and, importantly, did not show systematic species-driven differences **(Supplementary Figure 4A)**. Normalized enrichment scores for differentially expressed gene sets were in concordance with up-/downregulation of the tested sets across all studies, with generally high enrichment scores for upregulated gene sets and low enrichment scores for downregulated gene sets, supporting the existence of a shared core inflammation program. Of note, downregulation of genes was more variable across studies and hence less informative **(Supplementary Figure 4B)**. Pearson correlation coefficients of log_2_ fold change values showed strong positive skewing, again pointing towards a core inflammatory response across conditions and species **(Supplementary Figure 4C)**.

As a third analytical approach, we performed a WGCNA(39) that identified four modules with significant enrichment for core inflammatory response genes (Fisher’s exact test, *P*_*adj*_ < 0.05). Gene expression within those four modules increased in inflammation and contained several members of the core inflammation program **(Supplementary Figure 5)**.

On the level of individual samples, we could confirm that the group of 179 core inflammation genes had either weak or absent expression in healthy neutrophils and were induced in inflamed neutrophils **(Figure 3D)**.

Gene set enrichment analysis identified a conserved enrichment of pathways related to apoptosis, inflammatory response, IL-2 and IL-6 signaling, IFN-γ response, TNF signaling via NFKB and KRAS signaling **(Figure 3E)**.

Taken together, this integrative analysis of resting and activated neutrophils nominates a core inflammation program in neutrophils which is shared across inflammatory conditions and across species.

### The core inflammation program shows conserved transcriptional regulation across species

To identify putative regulators of neutrophil activation in inflammation, we applied transcription factor (TF) enrichment analysis individually to up- and downregulated genes in each study. TF enrichment across murine and human inflamed neutrophils was highly consistent in TFs with decreasing **(Figure 4A)** and increasing **(Figure 4B)** activity.

**Figure 4.**
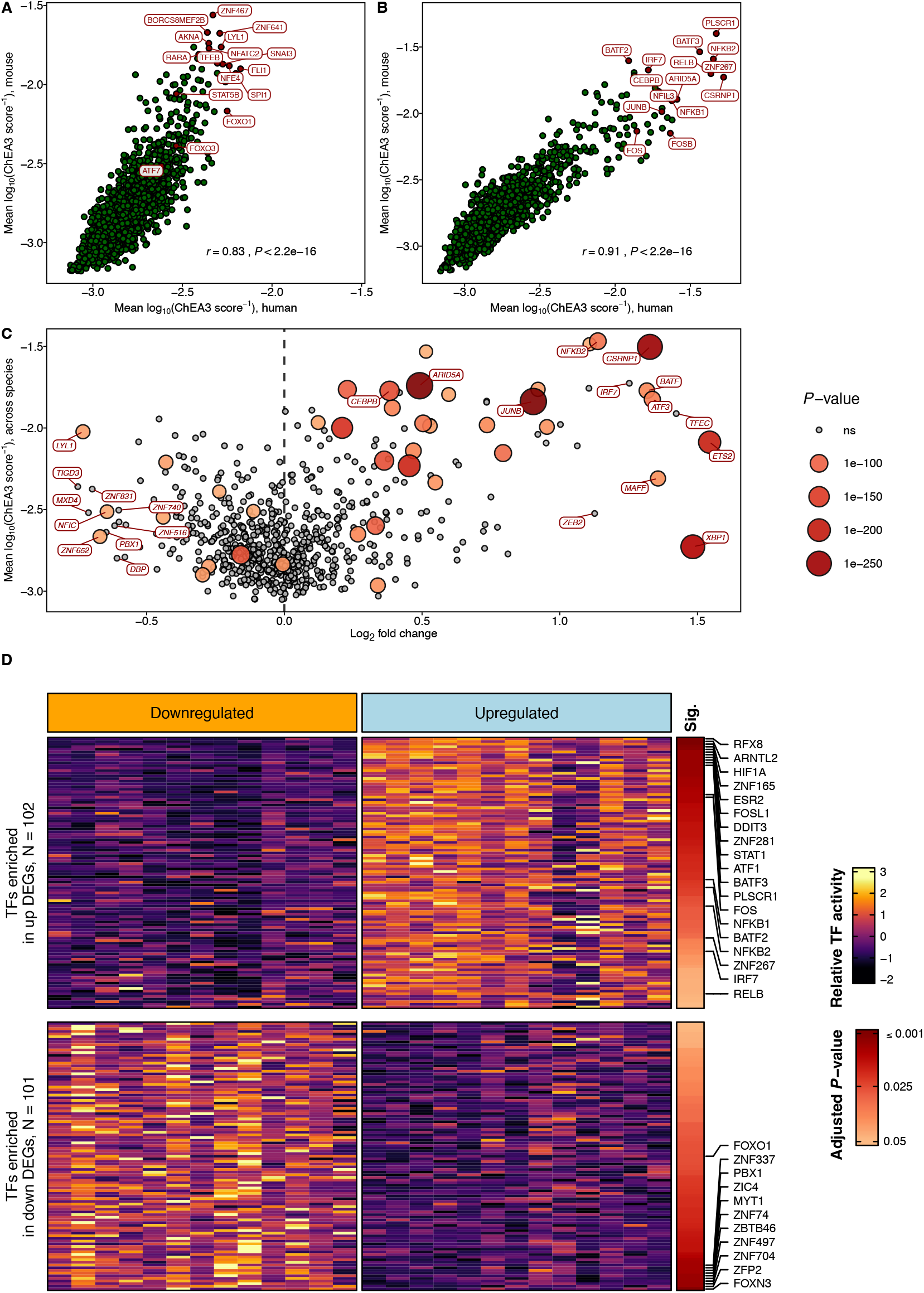
Transcription factor expression and regulatory activity in inflamed neutrophils is conserved across humans and mice. **(A)** Enrichment of transcription factors in genes associated with resting neutrophils is highly concordant between species. N = 250 genes with a negative log_2_(FC) in inflammation were sorted in ascending order of adjusted *P*-values for each comparison, and regulatory activity was derived as outlined in Methods. For each transcription factor, mean ChEA3-derived activity across human (x) and mouse (y) comparisons is shown; high values indicate high predicted activity. Top 10 transcription factors with the highest predicted activity as well as FOXO3, FOXO1, TFEB, RARA, STAT5B and ATF7 are highlighted. **(B)** Enrichment of transcription factors in genes associated with inflamed neutrophils is highly concordant between species. For each transcription factor, mean ChEA3-derived activity across human (x) and mouse (y) comparisons is shown; high values indicate high predicted activity. Top 10 transcription factors with the highest predicted activity as well as NFIL3, FOSB, FOS, CEBPB, and JUNB are highlighted. **(C)** A restricted set of transcription factors show both increased regulatory activity and expression in inflamed neutrophils. Shown is the mean log_2_ fold change of gene expressions across all comparisons (x; as in Figure 3B) versus the regulatory activity (inverse logarithm of ChEA3-Scores per species) (y). Only transcription factors where the respective gene had an assigned *P*-value in Fisher’s combined test (see Methods) are shown (N = 680). Colors and sizes indicate −log_10_(Fisher’s combined test). Genes encoding transcription factors with a mean log2 fold change ≥ 0 were merged with transcription factor scores that were derived from the upregulated score set (as in B), and genes with a mean log_2_ fold change < 0 were merged with transcription factor scores that were derived from the downregulated score set (as in A). Labeled are the genes encoding transcription factors with the 10 highest absolute log_2_ fold changes per direction which fall under the adjusted Fisher’s combined *P*-rank cutoff of 500 genes, as well as the 5 transcription factors with the lowest P-values and CEBPB. **(D)** Transcription factor enrichment in down- and upregulated genes in inflammation is shared across studies. Enrichment of transcription factors was calculated based on the top 250 down-(left) or upregulated (right) genes in each independent comparison in our differential expression analysis. The inferred activity of transcription factors (rows) is shown scaled across gene sets. Adjusted *P*-values from a t-test between inferred activity in down-vs. upregulated gene sets for each transcription factor are shown in the “Sig.” column. Transcription factors with the 10 lowest *P*-values per direction as well as those labeled in A-B were labeled.

Transcription factors that we found to be enriched in genes expressed in resting neutrophils include AKNA, PU.1 (encoded by *SPI1*), FOXO3, FOXO1, TFEB, RARA and STAT5B **(Figure 4A)**. Transcription factors that we found to be enriched in genes associated with inflamed neutrophils included CSRNP1, PLSCR1, FOS, FOSB, the NF-κB components NFKB1/NFKB2, the emergency granulopoiesis transcription factor CEBPB and JUNB **(Figure 4B)**.

To reduce this selection of transcription factors to those with the highest changes in inflammation, we compared predicted regulatory activity and actual transcription factor expression in inflammation. This analysis highlighted that CSRNP1, JUNB, CEBPB, XBP1, and ETS2 were strongly upregulated in inflamed neutrophils while also displaying strongly increased regulatory activity **(Figure 4C)**.

On the level of individual studies, we also found high consistency in the transcription factors predicted to be enriched in genes upregulated and downregulated in activated neutrophils **(Figure 4D)**. These results were consistent with an independent enrichment analysis performed separately for each species **(Supplementary Figure 6)**.

### Migration into tissue and activation significantly enhance chromatin accessibility and expression of core inflammation genes

If genes in the core inflammation program are predisposed to be upregulated, then chromatin accessibility for these genes should increase upon neutrophil maturation, migration into tissues and exposure to inflammatory stimuli.

To test this hypothesis, we analyzed ATAC-sequencing data derived from an air pouch model of acute inflammation, where neutrophils first migrate into a sterile membrane in the skin before being exposed to zymosan in the air pouch (36). Of the 179 core inflammation program genes, 29 displayed increasing accessibility in blood vs. bone marrow, compared to only 10 genes with decreased accessibility **(Figure 5A)**. Neutrophils which had transmigrated from blood into the membrane displayed enhanced accessibility of 78 genes. This increase was significantly (*P* = 5.1 × 10 ^−9^) higher than the increase of 29 genes for neutrophils in blood compared to bone marrow.

**Figure 5.**
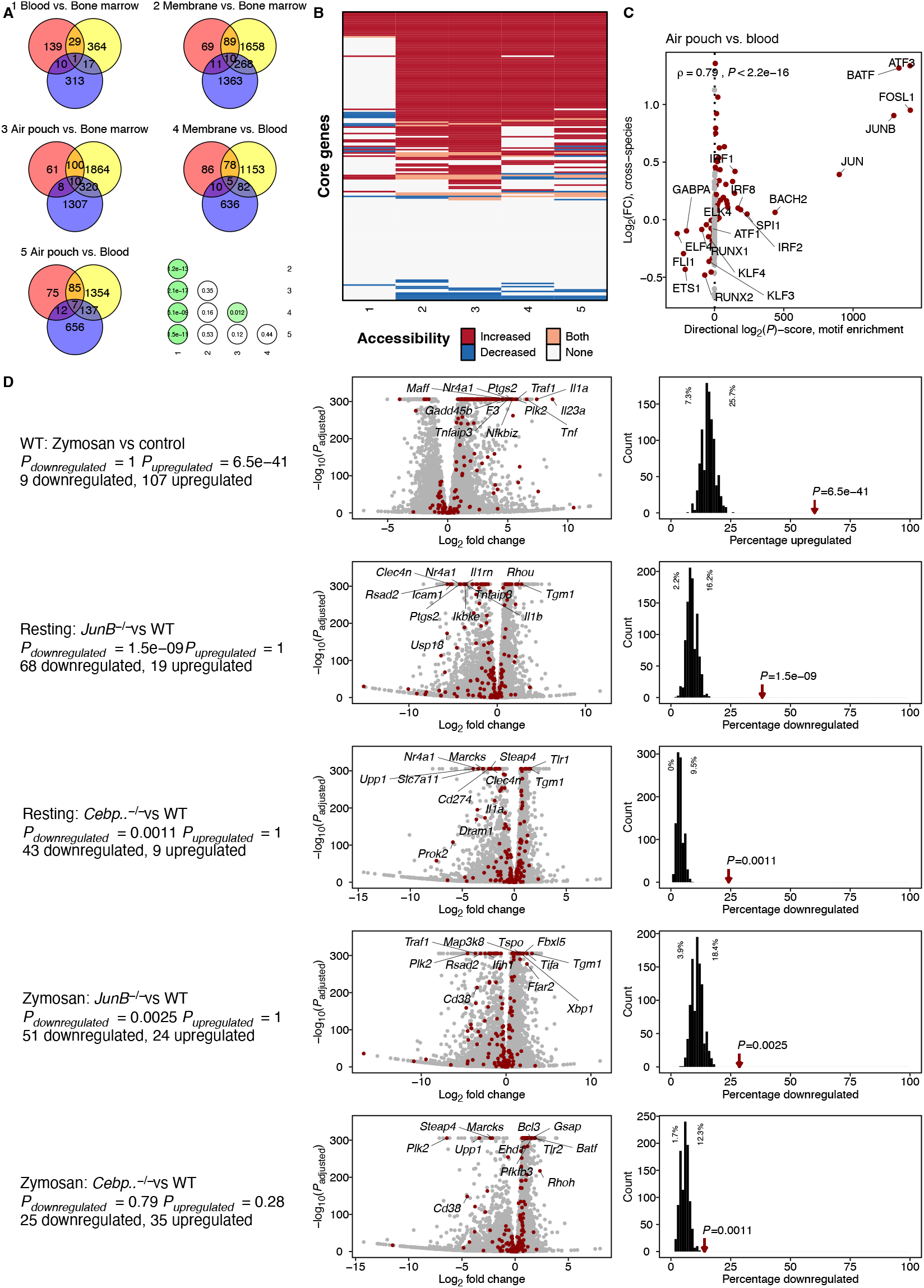
Genes in the core inflammation program are predisposed to be upregulated with maturation and activation. RNA-sequencing data and ATAC-sequencing data were retrieved from NCBI Gene Expression Omnibus (GSE161765). For A–C, we analyzed ATAC-sequencing results of the air pouch model (35, 47). C57BL/6J male mice received a dorsal air pouch and were challenged with zymosan into the pouch. Then, ATAC-seq was performed on neutrophils isolated from bone marrow, blood, membrane and exudate. For D, we restricted the analysis to HoxB8 cells that were treated as described (35). HoxB8 C57BL/6N myeloid progenitors (WT and CRISPR-Cas9-mediated targeted knockout cell lines) were differentiated to neutrophils for 5 days and were then challenged with zymosan for 2 hours (versus control). **(A)** Members of the core inflammation program show increasing chromatin accessibility in blood vs. bone marrow and in membrane and air pouch vs. bone marrow or blood. Venn diagrams derived from ATAC-sequencing data. Genes part of the inflammatory response program were compared with the list of genes that showed increased as well as decreased accessibility in the depicted comparisons. Bottom-right: Fisher’s exact test based on the number of core inflammatory genes with increased accessibility versus the number which did not. Shown are FDR-adjusted *P*-values. **(B)** Members of the core inflammation program showing increasing chromatin accessibility are shared across comparisons. Heatmap of core inflammation program gene accessibility for each comparison from (A). Rows were ordered by a nested decreasing rank of peaks associated with an increase, with both, with none and with a decrease. **(C)** Only a small subset of transcription factors show increased motif enrichment and increased gene expression in inflamed neutrophils. Motif enrichment analysis was performed using HOMER (see Methods) and are compared to the mean log_2_ fold change across species. ρ indicates Spearman’s rank correlation coefficient. **(D)** Core inflammation genes show preferential upregulation compared to random genes in zymosan-stimulated HoxB8 cells and are preferentially downregulated after JunB and Cebpb knockout. Comparison of core inflammation genes with HoxB8 neutrophil RNA-Seq data (from Khoyratty et al. (35)) Each tested comparison occupies one row. Experimental conditions for each comparison and respective statistics are shown in the left column. Volcano plots for each comparison are shown in the middle column. Core inflammation members are highlighted in red. Members with the highest combined significance and effect sizes are labeled. The right column contains histograms showing the percentage of random background genes (equally sized gene sets, 1000 simulations) up/downregulated (see x-axes) in each comparison. The red arrow indicates the observed percentage for core inflammation program members and is annotated with the respective overrepresentation *P*-value.

Similar skewing towards enhanced accessibility of core inflammation genes was also observed in the comparisons of membrane vs. bone marrow (89 up, *P* = 1.2 × 10 ^−13^), inflamed air pouch vs. blood (85 up, *P* = 1.5 × 10 ^−11^) and inflamed air pouch vs. bone marrow (100 up, *P* = 2.1 × 10 ^−17^), all compared to blood vs. bone marrow. **(Figure 5A)**. Furthermore, the genes with increased and decreased accessibility were highly consistent across the comparisons **(Figure 5B)**.

After finding that core inflammation genes have increased chromatin accessibility even before onset of inflammation, we searched for potential driver transcription factors displaying increasing expression and regulatory activity in inflammation. The regulatory activity was derived using HOMER motif enrichment analysis. Comparing motif enrichment with actual expression change in air pouch vs. blood, we observed an increase in both measures for a remarkably restricted set of transcription factors, namely ATF3, BATF, FOSL1, JUNB and JUN **(Figure 5C)**.

We next investigated whether the core inflammation program represents a group of genes from which neutrophils preferentially draw upon exposure to inflammatory stimuli. If this was the case, then it should be more likely for core inflammation genes to be upregulated in inflammation compared to all other genes. We analyzed RNA-seq data from differentiated HoxB8 neutrophils stimulated with or without zymosan for 2 hours (36). We observed that in activated neutrophils, a significantly higher proportion of core inflammation genes (107/179 = 60 %) was upregulated than expected by chance (15 – 46 genes in 1,000 simulations using random background genes; P_overrepresentation_ = 6.5 × 10 ^−41^) **(Figure 5D)**.

To assess the impact of two transcription factors identified in our enrichment analysis on expression of core inflammation program genes, we repeated the same analysis in differentiated HoxB8 neutrophils carrying a genetic knockout of either *JunB* or *Cebpβ*. CEBPB is a key transcription factor mediating emergency granulopoiesis (40) and showed upregulation in inflamed neutrophils as well as increased regulatory activity. In addition, JUNB, which plays an important role in the inflammatory response of neutrophils (9, 36), also was had increased motif enrichment in the air pouch vs. blood comparision.

Based on these analyses, we expected a modest reduction in expression of core inflammation genes in *Cebpβ*^−/−^ cells and a stronger reduction in *JunB*^−/−^ cells. Indeed, this was the case: In a direct comparison of resting knockout (*JunB*^−/−^ and *Cebpβ*^−/−^) versus wildtype cells, we observed a significantly stronger downregulation of the core inflammation program in *JunB*^−/−^ cells (69 genes; *P* = 1.5 × 10 ^−9^) than in *Cebpβ*^−/−^ cells (43 genes; *P* = 0.0011) **(Figure 5D)**.

Comparing zymosan-stimulated knockout cells versus wildtype cells, we again saw a significant downregulation of core inflammation genes in the *JunB*^−/−^ condition (51 genes; *P* = 0.0025) but not in the *Cebpβ*^−/−^ condition (25 genes; P = 0.79) **(Figure 5D)**.

Together, these results indicate that maturation and migration into an inflamed tissue site predispose neutrophils to upregulate genes of the core inflammation program and that knockout of *Cebpβ* and especially *JunB* leads to a weaker induction of core inflammation genes compared to WT cells.

### Members of the core inflammation program can be validated on the protein level in activated human and murine neutrophils

To validate members of the core inflammation program experimentally, we filtered the list of genes by surface proteins, yielding 36 markers **(Figure 6A)** (41). Based on antibody availability, we developed a flow cytometry panel including canonical lineage markers (human: CD15, mouse: Ly6G) and five proteins predicted to be part of the core inflammation program: CD14, CD69, CD40, CD274 (PD-L1) and IL-4R **(Supplementary Tables 3 and 4)**.

**Figure 6.**
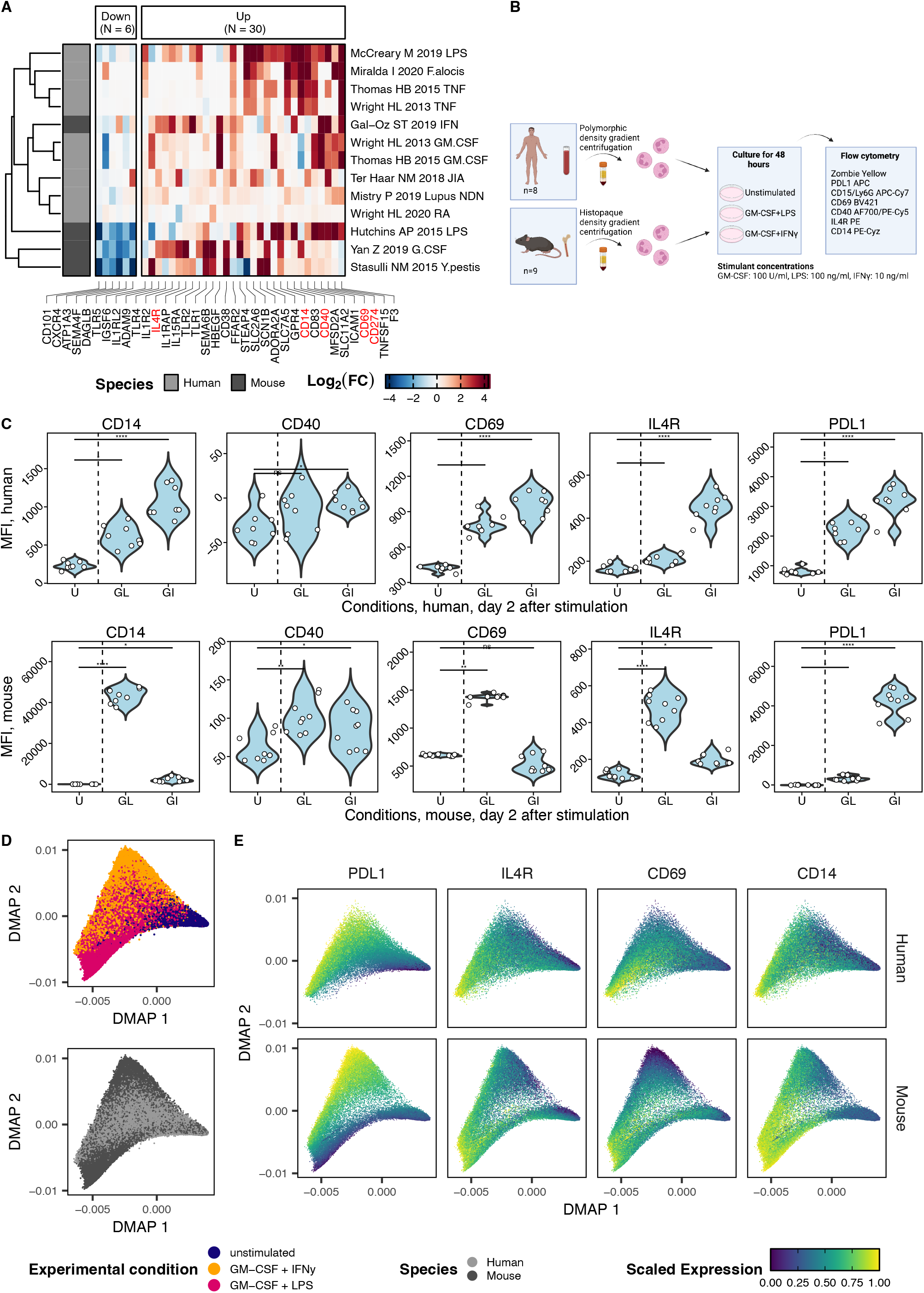
Experimental validation of the core inflammation program on the protein level. **(A)** Inflammation-specific protein-coding genes were identified by filtering the genes depicted in Figure 3C for the surfaceome as previously described (41). Genes that encode proteins selected for validation (based on antibody availability and panel design) are labeled in red. **(B)** Experimental overview. Human and murine neutrophils were isolated from peripheral blood or bone marrow, respectively, cultured 48 hours and analyzed in flow cytometry. **(C)** Flow cytometry analysis of resting and activated murine and human neutrophils. The gating strategy is depicted in Supplementary Figure 8. Significance indicates adjusted P-values of a Dunn’s test that followed a Kruskal-Wallis H test (significant for all markers). **(D)** Top, diffusion map embedding of neutrophils from humans and mice cultured for 48 hours with or without GM-CSF + LPS and GM-CS F + IFN-γ. Diffusion map embedding calculated based on CD69, CD14, IL-4R and PD-L1. Bottom, diffusion map embedding of human and murine neutrophils, colored by species. **(E)** Diffusion map embedding colored by marker expression highlights a continuum driven by increasing

We isolated human neutrophils from peripheral blood and murine neutrophils from bone marrow and cultured them over 48 hours with or without the addition of GM-CSF + LPS and GM-CSF + IFN-γ **(Figure 6B)**.

Aging in cell culture without activation led to a significant upregulation of CD69 (in human) and to a significant downregulation of CD14 and IL4R both in human and in mouse **(Supplementary Figure 7)**. Compared to unstimulated cells, activated murine neutrophils significantly upregulated the predicted core inflammation program markers CD14, CD40, CD69, PD-L1 and IL-4R in the condition containing LPS and all but CD69 in the condition containing IFN-γ **(Figure 6C)**. Human neutrophils displayed a highly concordant increase in those markers. CD69 and PD-L1 increased with similar magnitude, while upregulation of CD14 was stronger in murine neutrophils compared to human neutrophils. In human neutrophils, upregulation of CD40 was restricted to a small (∼ 2%) population of neutrophils but reached significance on the bulk level for GM-CSF + IFN-γ stimulation **(Figure 6C)**.

Differences were also noticeable between inflammatory conditions. In murine neutrophils, the combination of GM-CSF and LPS led to a stronger increase in expression of CD14, CD69, IL-4R and CD40 compared to GM-CSF and IFN-γ. The reverse was true for PD-L1, which is driven substantially by IFN-γ signaling (8). In mice, IFN-γ stimulation reduced CD69 expression, while LPS increased it. In human neutrophils, the combination of GM-CSF and IFN-γ lead to stronger increases in CD14, CD69, IL-4R and PD-L1 than the combination of GM-CSF and LPS.

A combined diffusion map analysis revealed a high degree of overlap between murine and human neutrophils, while cell distribution was driven predominantly by experimental condition **(Figure 6D)**. Correspondingly, activated neutrophils of both species displayed a continuous upregulation of the inflammatory response markers **(Figure 6E)**.

These findings confirm the predicted activation markers, further substantiating the conservation of inflammatory response programs in neutrophils while also revealing differences between species and inflammatory conditions.

### Neutrophil origin and inflammatory condition influence expression of the core inflammation program

Neutrophil heterogeneity is influenced by the tissue microenvironment (6, 9). To evaluate the impact of tissue origin on the phenotype of neutrophils in inflammation, we performed stimulation experiments with paired leukocyte preparations from blood, bone marrow and spleen of wild-type BL6 mice. In a principal component analysis of flow cytometry data, resting neutrophils clustered closely together but each tissue remained distinguishable based on subtle baseline expression differences in IL-4R, CD69 and CD40 **(Figure 7A)**. Inflamed neutrophils deviated markedly from their resting counterparts and reached distinct states as a function of tissue and inflammatory condition **(Figure 7A)**.

**Figure 7.**
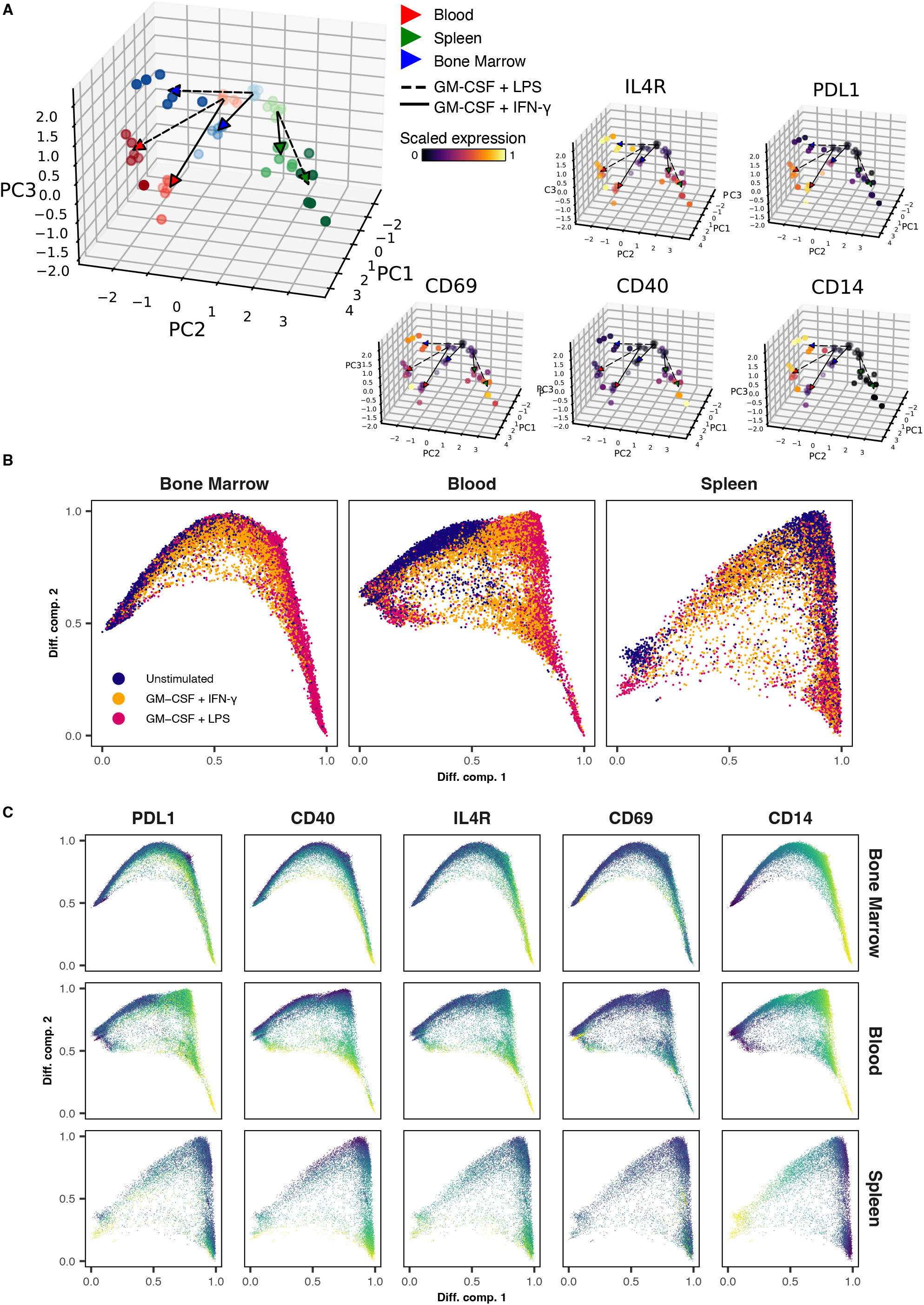
Neutrophil origin and cytokine stimulation determines manifestation of the core inflammation program. **(A)** Left: Principal component analysis performed on the median marker expressions per sample reveals distinct phenotypes of neutrophils driven by organ of isolation (color coded) as well as cytokine stimulation (line-segment coded). Right: PCA plots colored by scaled expression for each of the quantified markers. **(B)** Diffusion Map embedding of neutrophils from bone marrow, blood and spleen cultured for 8 hours with or without GM-CSF + LPS and GM-CSF + IFN-γ. Embeddings were calculated based on the expression levels of CD69, CD14, IL-4R, PD-L1 and CD40. **(C)** Diffusion Map embedding of resting and cytokine-stimulated neutrophils (as in B) from bone marrow, blood and spleen, colored by marker expression.

Neutrophils from all tissues upregulated CD69 and IL-4R, suggesting that these markers can be utilized as neutrophil activation markers across a variety of conditions **(Figure 7A)**. In contrast, expression of CD40, CD14 and PD-L1 showed greater tissue dependence. CD40 (evident most prominently in murine neutrophils) was robustly upregulated in splenic neutrophils and less prominently in blood neutrophils. Conversely, CD14 and PD-L1 expression was inducible to a greater extent in blood neutrophils and bone marrow neutrophils but less in splenic neutrophils.

We also noted differences related to activating stimuli, for example, through more prominent PD-L1 induction by IFN-γ compared to LPS. Single-cell analysis highlighted a continuum of states in all organs **(Figure 7B)**, driven by increasing expression of the core inflammation markers **(Figure 7C)**. Importantly, the core program was already inducible in bone marrow neutrophils, supporting the validity of *in vitro* and adoptive transfer experiments performed with bone marrow neutrophils.

Together, these findings experimentally validate the core inflammation program in neutrophils from different tissues and highlight that organ and stimulus further influence the resulting neutrophil phenotype.

## DISCUSSION

Neutrophils are important mediators of immune defense and protagonists in immune-mediated diseases. Murine and human neutrophils differ in morphology, frequency in blood (humans ∼ 50– 70%, mice ∼ 10–25%) and expression of marker proteins. For example, murine neutrophils are defined by surface expression of Ly6G, not present in the human genome, whereas murine neutrophils lack expression of defensins (42).

Both in humans and mice, neutrophils are phenotypically heterogeneous across different tissues and inflammatory conditions (37, 43, 44). Recent studies suggest that neutrophil heterogeneity in homeostasis is driven by a chronological sequence of maturation and activation termed neutrotime, whereas the combination of aging, tissue factors, environmental features and inflammatory signals promote their polarization towards distinct states (6, 7, 9).

While the neutrotime signature can be detected in both species and this overarching principle of neutrophil ontogeny is likely conserved across humans and mice, it is poorly understood which features of the neutrophil inflammatory response are shared across species. Furthermore, it is unclear which aspects of the neutrophil inflammatory response reflect a general inflammatory response program shared across multiple inflammatory conditions and which features are highly specific to certain triggers or sites of inflammation.

To address these gaps in knowledge, we performed an integrative analysis of 76 leukocyte samples from the Haemopedia atlas, 195 additional samples from humans and mice obtained through the ENA for the analysis of resting and inflamed neutrophils, 18 HoxB8-derived samples for the analysis of gene knockouts, and murine ATAC-sequencing data from different tissues. We integrated these datasets using high-confidence orthologs and validated our computational approach by comparing gene expression conservation across six immune cell lineages: T cells, B cells, monocytes, dendritic cells, NK cells and neutrophils. Expression of lineage-specific genes was generally well-conserved across humans and mice. Intriguingly, neutrophils displayed both the greatest number of lineage-specific genes and the highest correlation of gene expression between mice and humans, suggesting a higher degree of conservation in this phagocytic cell compared to other lineages.

We also evaluated eleven studies of experimental inflammation. While different conditions induced highly heterogeneous responses in neutrophils, our combined analysis allowed us to predict a core inflammation program conserved across murine and human neutrophils. While studies display variability with respect to genes upregulated in activated neutrophils, a core inflammation program emerged from which genes are preferentially upregulated. The robustness of this program was underscored by the high concordance between the gene set derived from Fisher’s combined test and complementary approaches based on a linear mixed model as well as weighted correlation network analysis (WGCNA; (39, 45, 46)). The conservation of a small set of transcription factors predicted to regulate a broad variety of conditions across humans and mice highlights the conserved nature of gene expression in neutrophils.

To validate the predicted core inflammation program in different models, we analyzed differential gene accessibility in published ATAC-sequencing data from a murine air pouch model of inflammation. We found a significant proportion of core inflammation program genes to be more accessible with maturation and in the pro-inflammatory conditions, highlighting the predictive value of the program in a method not used in the generation of the program.

HoxB8-derived neutrophils are a powerful tool to model neutrophil function. We assessed the differential expression of zymosan-activated HoxB8-derived neutrophils versus control, showing a significant overrepresentation of core inflammation genes in activated neutrophils. The core inflammation program was reduced in resting cells carrying a knockout of key regulators of this program (*JunB*^-/-^ and *Cebpβ*^-/-^). Core inflammation genes were also significantly less upregulated in zymosan stimulated *JunB*^-/-^ cells, indicating an impaired neutrophil inflammatory response in these cell lines. Concordant with previous reports of a more limited impact of the *Cebpβ* knockout on inflammatory neutrophil functions compared to the *JunB* knockout (36), the underrepresentation of core inflammation genes was nonsignificant in our analysis.

Finally, we validated key components of the predicted core inflammation program experimentally. Using primary human and murine neutrophils, we showed that the surface proteins CD14, CD69, IL-4R, CD40 and PD-L1 are induced by *in vitro* cytokine stimulation, and this upregulation is observable in both species, although CD40 was restricted to a small subset of neutrophils in humans. This finding further underlines the conserved character of the inflammation program as presented in this study. Interestingly, while neutrophils from different murine tissues upregulated the inflammatory response markers, the magnitude of upregulation differed across bone marrow, spleen and blood, suggesting that the tissue origin of neutrophils is an important consideration in experimental studies.

Our study was limited to bulk RNA-sequencing samples, since a similar analysis using single-cell studies requires datasets that are only now beginning to emerge. To circumvent potential batch effects, we focused our analysis on studies with internal controls of resting neutrophils, excluding other potentially interesting studies containing only neutrophils harvested from inflamed sites.

Nevertheless, our combined analysis of 11 human and murine neutrophil transcriptomic datasets identified a largely conserved transcriptomic landscape across species, supporting the use of murine neutrophils to illuminate human biology. We furthermore predicted and experimentally confirmed the existence of a core inflammation program conserved across human and murine inflamed neutrophils. Going forward, genetic perturbations and pharmacological interventions to interfere with pathologic neutrophil activation will be particularly informative if focused on this conserved program. The systems biology approach presented here can be transferred to other cell types and organisms to facilitate further studies comparing gene expression across species.

## METHODS

Methods are available in the supplementary information.

## ACKNOWLEDGMENTS

This work was supported by MD fellowships from the Boehringer Ingelheim Fonds (to N.S.H., F.A.R.), a MD/PhD fellowship by the Medical Faculty of Heidelberg (to F.A.R., T.E.). R.G-B. was supported by the state of Baden-Wuerttemberg within the Centers for Personalized Medicine Baden-Wuerttemberg (ZPM), a physician-scientist development grant from the Medical Faculty Heidelberg, and a research grant from the German Society for Rheumatology (DGRh). Figures **1A** and **6B** were created with BioRender.com. The authors acknowledge support by the state of Baden-Württemberg through bwHPC and the German Research Foundation (DFG) through grant INST 35/1597-1 FUGG.

## AUTHORSHIP

Contribution: N.S.H. and F.A.R. conceptualized and designed the study, performed computational and statistical analysis, provided conceptual input in experimental planning, analyzed experiments, conceptualized the core inflammation program and wrote the manuscript. T.E. planned experiments, processed human and murine samples, performed and analyzed flow cytometry and wrote the manuscript. H.-M.L., C.M.-T., P.A.N., G.W. guided data analysis, interpretation and validation. R.G.-B. conceptualized and designed the study, supervised the entire project and wrote the manuscript.

## CONFLICT-OF-INTEREST DISCLOSURE

The authors declare no competing financial interests.

### Patient consent for publication

Not required.

### Ethics approval

EDTA-anticoagulated blood from healthy donors was collected under IRB-approved protocols Heidelberg S-285/2015.

## CODE AND DATA AVAILABILITY

Code for all analyses that are described in this manuscript will be made available in a public repository upon publication. RNA-Sequencing data is available from public databases (see Supplementary Information).

