## Supplementary Information for "Human and murine neutrophils share core transcriptional programs in both homeostatic and inflamed contexts"

### SUPPLEMENTARY METHODS

#### Datasets

For all analyses, we used the following datasets:

##### *RNA-Sequencing*

Datasets of interest were identified through a literature search on PubMed and the NCBI Gene Expression Omnibus. In total, 262 publicly available RNA sequencing samples from 24 studies were included.

- Lineage atlas dataset: 76 samples, curated subset of the Haemopedia RNA-Seq atlas available through the SRA
  - Human: 40 samples
    - B cells 8 samples
    - Dendritic cells 7 samples
    - Monocytes 7 samples
    - Neutrophils 3 samples
    - NK cells 5 samples
    - T cells 10 samples
  - Mouse: 36 samples
    - B cells 2 samples
    - Dendritic cells 4 samples
    - Monocytes 6 samples
    - Neutrophils 6 samples
    - NK cells 2 samples
    - T cells 16 samples
- Neutrophil dataset: 195 samples (including the 9 haemopedia neutrophil samples), curated list of studies available through the SRA. All of the studies below were selected only to contain neutrophils. The total amount of samples for each study is shown; studies that were selected for inflammatory differential expression analysis are highlighted in bold. Subsets of samples from studies not selected for differential expression analysis have been used in analyses focusing on healthy control samples (Fig 2).
  - Human: 136 samples from 13 studies
    - Adrover JM 2020 FALSE 6 samples
    - Catapano M 2019 FALSE 11 samples
    - de Graaf CA 2018 FALSE 3 samples (haemopedia atlas neutrophil samples)
    - Franco LM 2019 FALSE 2 samples
    - Grabowski P 2019 FALSE 22 samples
    - Vecchio F 2018 FALSE 8 samples
    - **McCreary M 2019 TRUE 10 samples**
    - **Miralda I 2020 TRUE 16 samples**
    - **Mistry P 2019 TRUE 28 samples**
    - **Ter Haar NM 2018 TRUE 6 samples**
    - **Thomas HB 2015 TRUE 12 samples**
    - **Wright HL 2013 TRUE 6 samples**
    - **Wright HL 2020 TRUE 6 samples**
  - Mouse: 59 samples from 11 studies
    - Bhalla M 2021 FALSE 7 samples
    - Casulli J 2019 FALSE 3 samples

- Coffelt SB 2014 FALSE 4 samples
- de Graaf CA 2018 FALSE 6 samples (haemopedia atlas neutrophil samples)
- Germann M 2020 FALSE 4 samples
- Hsu BE 2019 FALSE 4 samples
- Zhu YP 2018 FALSE 3 samples
- **Gal-Oz ST 2019 TRUE 12 samples**
- **Hutchins AP 2015 TRUE 4 samples**
- **Stasulli NM 2015 TRUE 6 samples**
- **Yan Z 2019 TRUE 6 samples**

***RNA-Seq of HoxB8 cells***

- Khoyratty TE 2021 18 samples

***ATAC-Seq***

- Khoyratty TE 2021 5 peak annotations

***Flow cytometry***

- This study samples from 8 human donors and 9 mice

**Data retrieval and processing**

We downloaded raw sequencing reads for the selected studies to the MLS&WISO bwForCluster using release 1.5 of the nf-core(1) fetchngs pipeline and quantified them using release 3.6 of the nf-core rnaseq pipeline. The pipelines were launched using nextflow(2) version 22.04.0. To ensure high reproducibility, all pipeline processes were run inside singularity (version 3.9.2) containers. We mapped all downloaded samples using salmon(3) version 1.5.2 with the parameters libType set to 'A' and indexing the reference genomes with 21 base k-mers (**Supplementary Table 5**). Quantified transcripts were summarized to the gene level using bioconductor-tximeta(4) version 1.8.0. All human samples were mapped to the GRCh38 genome. All mouse samples were mapped to the GRCm39 genome unless stated otherwise. Author-supplied metadata was queried using GEOquery (v2.64.0; (5)) and integrated manually to ensure consistency across studies **(Supplementary Table 1)**. R (v4.2.0) was used for downstream analyses. Bioconductor (v3.15) and additional packages were used for downstream analyses and visualizations (6-9).

**Orthology analyses and mapping**

For downstream analyses, genes were mapped using ENSEMBL Version 107 (10). We restricted all our composite cross-species analyses to protein-coding genes with a high-confidence orthology relationship and available gene symbols in both species. Murine and human gene expression datasets were combined based on these orthologs.

**Identification of lineage-associated genes**

Lineage-associated genes were identified using a linear model based differential expression test, implemented in limma (v3.52.0; (11)) and edgeR (v3.38.0; (12-14)). Differential expression testing was restricted to protein-coding genes that could be assigned high-confidence orthologs between human and murine samples. We constructed a cross-species count matrix based on those mappings and referred to each mapped gene by its human gene symbol. Counts were filtered using edgeR's filterByExpr filtering approach. We applied TMM normalization to account for differences in library composition. We then transformed counts to $\log_2$(CPM) values and estimated weights for each observation using voom. We applied limma to fit a linear model to our data and calculated differential expression for a given lineage against all remaining lineages. Lineage-associated genes were defined as genes that were differentially expressed in each lineage against all other lineages at a Benjamini-Hochberg corrected *P*-value of ≤ 0.05 and a $\log_2$ FC > 0. Genes were ranked according to their F statistic, and up to 200 genes were selected per lineage.

**Lineage PCA, Correlation analysis** and **clustering**

We used these balanced lineage-associated gene sets to perform PCA as well as correlation and clustering analysis on all samples. Human and murine samples were combined as described above. To emphasize our focus on comparisons between lineages, we mean-centered $\log_2$(CPM) for each species prior to combining the count matrices. A PCA was computed for all integrated samples, taking the concatenated lineage-associated gene sets as input features. Correlation of expression analyses was performed based on the same features, calculating Pearson's *r* correlation coefficient for each inter-sample combination. We subsequently performed a hierarchical clustering analysis on the obtained correlation coefficients.

**Comparison of neutrophil lineage gene expression profiles in resting neutrophils**

To compare expression patterns of neutrophil lineage-associated genes in resting human and murine neutrophils, we first defined lineage-associated genes for human and murine samples separately. We defined those genes as lineage associated that were upregulated (Benjamini-Hochberg corrected *P*-value of ≤ 0.05 and a $\log_2$ FC > 1) in neutrophils against all other lineages. We next mapped those gene sets to their human and murine counterparts, considering only high confidence one-to-one, one-to-many, and many-to-many orthology relationships. Based on those mappings, we merged all genes detected as lineage-specific in either of the considered species. We also included genes detected as lineage-specific in either species but could not be mapped to a high-confidence ortholog. The obtained genes were subset only to include genes that showed evidence of expression in the inflammatory dataset.

Taking the computed mappings and $\log_2$(TPM+1) expression values of mapped gene-gene pairs, we tested for differential expression of those pairs between species using a linear mixed model (lme4 v1.1-29; (15)) accounting for study-related batch effects by including the study annotation as a random effect:

Full model: $\log_2$(TPM+1) ~ species + (1|study)

Null model: $\log_2$(TPM+1) ~ (1|study)

*P*-values were computed by performing a likelihood ratio test between these models. We subsequently adjusted those values using the Benjamini-Hochberg correction method based on the total number of tested gene-gene pairs (genes that appeared as lineage-specific in either species were expressed in the inflammatory dataset and could be mapped to one or more counterparts with high confidence).

Using the average expression of mapped gene-gene pairs and differential expression P-value, we defined 5 different expression profiles: Genes that showed high (> $95^{th}$ percentile of all genes that were detected as lineage-specific in either species) average expression levels in both species and did not exhibit differential expression between species (Benjamini-Hochberg corrected P-value > 0.05, absolute beta < 1). Additionally, we defined 4 divergent clusters of genes that had high expression levels in only one of both species and showed evidence of differential expression (Benjamini-Hochberg corrected P-value < 0.05, absolute beta ≥ 1) or were abundantly expressed but could not be assigned an orthologous gene in the other species respectively.

#### Differential expression testing

We performed differential expression analyses between inflamed and resting conditions on a total of 112 samples from N = 11 (human: 7, mouse: 4) studies. To account for potential batch effects between studies, we used DESeq2 (v1.36.0; (16)) in each of the studies individually to identify differentially expressed genes in inflamed compared to healthy control samples. Each study's gene list was pre-filtered to only include genes with counts > 1 in at least 1 sample before differential expression analysis, based on the negative binomial distribution. To remove noise while preserving significant differences, $\log_2$ fold change results were then shrunk using the *apeglm* package (17). Differential gene expression results were additionally filtered through DESeq2's default independent filtering approach, as well as its count outlier filtering.

#### Identification of a core inflammation program

To assess which inflammation-driven changes in the neutrophil transcriptome are shared across conditions and conserved across species, we applied a Fisher's combined test to the adjusted *P*-values of each gene in each study, restricting the analysis to genes that passed our expression filter as well as DESeq2 filters in ≥ 80% of studies. This analysis provided a Benjamini-Hochberg-corrected composite *P*-value for all genes and allowed us to rank genes by their likelihood of dysregulation in inflammation. Additionally, we calculated a mean $\log_2$ fold change for each gene across all studies.

Based on a rank-*P*-value plot (**Figure 3B**), we determined a *P*-value cutoff at a rank equaling 500, corresponding to an adjusted Fisher $P \leq 6.164117 \times 10^{-40}$. Genes with an absolute $\log_2$ fold change greater than or equal to 0.5 and an adjusted *P*-value below our defined threshold were considered conserved in inflammation. We defined the upregulated subset ($\log_2$ FC ≥ 0.5) of those conserved genes as the core inflammation program.

#### Pathway enrichment analysis

Inflammatory pathway enrichment analysis was performed for each study individually using the fGSEA implementation of the Gene Set Enrichment Analysis method. For each study, the differential expression analysis results were ranked by $\log_2$ fold change. Enrichment was calculated for hallmark gene sets that were retrieved from the Molecular Signatures Database (v7.5.1; (18)).

#### Transcription Factor Enrichment Analysis

In order to identify regulators associated with genes induced or downregulated in inflammation, we used ChEA3 (21) with default settings as described (20), using the 250 most significantly up- and downregulated genes, respectively, for each condition, ranked by their adjusted *P*-value. We then calculated the arithmetic mean of the negative logarithms of the ChEA3 scores per species and transcription factor to compare average TF activity across species.

We used a paired t-test to assess significant differences between a TFs ChEA3-enrichment in up- against downregulated genes across all comparisons. Resulting P-values were corrected using a Benjamini-Hochberg correction for all tested TFs. We used these corrected P-values to determine if a TF was significantly more enriched in genes upregulated in inflammation or vice-versa. We subsequently inferred transcription factor activity using DoRothEA (v1.8.0; (22)) and decoupleR (v2.2.2; (23)), taking advantage of the species-specific transcription factor databases. Here, $\log_2$ fold change matrices per species served as input, leading to enrichment scores with their respective *P*-values. For downstream analyses, we calculated the mean enrichment scores per species and preserved the highest observed *P*-value for each transcription factor.

#### ATAC-sequencing analysis

We retrieved ATAC-sequencing data from mice that were subjected to the air pouch model of acute inflammation (GEO: GSE161765, mapped to the GRCm38 genome). Genes annotated based on differentially accessible peaks as defined in the study ($P_{adj} < 0.05$, fold change > 1.5) were compared with the conserved upregulated genes as defined in the core inflammation program. The ratio and number of core inflammation program genes that were associated with projected increased accessibility served as an input for pairwise Fisher's exact tests, *P*-values were adjusted using the Benjamini-Hochberg method.

#### RNA-sequencing analysis of zymosan-treated HoxB8 cells

We retrieved featureCounts (per ENSEMBL-ID) from HoxB8 cells that were subjected to differentiation and zymosan-treatment (GEO: GSE161765, mapped to the GRCm38 genome). Differential expression analysis was performed as described in the respective section above. We restricted the analysis on HoxB8-cells that were differentiated for 5 days and then compared (1) wildtype cells that were treated for 2 hours with zymosan (50 µg/ml) or DMSO (control), (2) resting stable knockout HoxB8 cell lines versus wildtype, (3) zymosan-treated stable knockout HoxB8 cell lines versus zymosan-treated wildtype HoxB8 cells. A significant up- or downregulation of core inflammation program genes was then assessed by performing pairwise overrepresentation analyses as explained above.

#### Experimental validation

The list of ranked conserved inflammatory response genes was filtered to include genes encoding surface proteins using the surfaceome resource (24). The remaining n=69 surface protein-encoding genes were then filtered by available human and mouse antibodies (BioLegend), and a panel consisting of CD14, CD69, CD40, IL-4R and PD-L1 was selected for validation.

**Supplementary Table 3. Human neutrophil flow cytometry panel**

| Marker | Channel | Clone | Vendor | Catalog # | dilution |
|---|---|---|---|---|---|
| **LIVE/DEAD** | Pacific Orange | N/A | BioLegend | 423103 | 1:300 |
| **CD15** | APC-Cy7 | W6D3 | BioLegend | 323047 | 1:100 |
| **CD69** | BV421 | FN50 | BioLegend | 310929 | 1:50 |
| **CD40** | Alexa Fluor 700 | 5C3 | BioLegend | 334327 | 1:50 |
| **CD14** | PE-Cy7 | M5E2 | BD Biosciences | 557742 | 1:100 |
| **IL4R** | PE | G077F6 | BioLegend | 355003 | 1:100 |
| **PD-L1** | APC | 29E2A3 | BioLegend | 329708 | 1:20 |

**Supplementary Table 4. Murine neutrophil flow cytometry panel**

| Marker | Channel | Clone | Vendor | Catalog # | dilution |
|---|---|---|---|---|---|
| **LIVE/DEAD** | Pacific Orange | N/A | BioLegend | 423103 | 1:300 |
| **Ly6G** | APC-Cy7 | 1A8 | BioLegend | 127623 | 1:100 |
| **CD69** | BV421 | H1.2F3 | BioLegend | 104527 | 1:100 |
| **CD40** | PE-Cy5 | 3/23 | BioLegend | 124617 | 1:100 |
| **CD14** | PE-Cy7 | Sa14-2 | BioLegend | 123315 | 1:100 |
| **IL4R** | PE | I015F8 | BioLegend | 144803 | 1:100 |
| **PD-L1** | APC | 10F.9G2 | BioLegend | 124311 | 1:100 |

#### Human samples

Research with healthy human participants followed the declaration of Helsinki. Peripheral blood of healthy donors was collected under an IRB-approved protocol (Heidelberg-S-272/2021). Neutrophils were isolated using density gradient centrifugation with Polymorphprep as previously described (25): 30 ml whole blood was layered onto 20 ml Polymorphprep (Progen #1114683) and centrifuged at 535 g for 35 min. The PBMC-containing layer was discarded by suction and neutrophils were recovered and subjected to hypotonic lysis using 0.2% NaCl. The cells were subsequently washed with cell culture medium (RPMI 1640 (Gibco #21875-034)) supplemented with 10% heat-inactivated FBS (PAN Biotech #3302/P101102) and 1% GlutaMAX (Gibco #35050-061) and seeded at 5 million cells per 6 wells in a total volume of 5 ml at a humidified atmosphere at 37°C with 5% $CO_2$. The cells were cultured over 48h either in the absence of cytokines (vehicle control), with GM-CSF + IFN-γ or GM-CSF + LPS. GM-CSF was used at a final concentration of 100 U/ml (R&D #215GM), IFN-γ at 10 ng/ml (BioLegend #570208) and LPS at 100 ng/ml (Invivogen #tlrl-3pelps). After 48 h, 1 million cells were collected and stained using the Zombie Yellow Fixable Viability Kit (BioLegend #423103) for live/dead discrimination, followed by an antibody panel **(Supplementary Table 3)** in 50 µl of FACS buffer (2% FBS, 5 mM EDTA and 0.1 sodium azide in PBS) for 25 min.

#### Murine samples

Experiments were conducted under the approval of the Animal Care Facility Heidelberg and the Animal welfare officers (approval #T66/21). Male and female C57BL/6J mice were sacrificed by cervical dislocation and bone marrow was extracted by flushing with RPMI. Neutrophils were enriched by density centrifugation using Histopaque 1077 (Sigma-Aldrich #10771) and Histopaque 1119 (Sigma-Aldrich #11191). Cells were recovered from the interphase of both Histopaque layers and centrifuged. Cells were washed with RPMI containing 10% FBS and 1% Glutamax and seeded at $10^6$ cells/ml in 48 well plates in a total volume of 500 µl. Murine GM-CSF (Peprotech #315-03, 100 U/ml), murine IFN-γ (Peprotech #315-05, 10 ng/ml) and LPS (Invivogen #tlrl-3pelps, 100 ng/ml) were added to the medium for 24 hours and 48 hours in combination as indicated in the respective figures. Cells cultured in the absence of cytokines were used as controls. After the indicated times, cells were collected and stained with the antibody panel (**Supplementary Table 4**) in 50 µl of FACS buffer containing 2% FBS, 5 mM EDTA and 0.1% sodium azide.

To assess neutrophils from different organs, mice were sacrificed by cardiac puncture under generalized anesthesia. Subsequently, the femora and tibiae were flushed with PBS to obtain bone marrow. Any remaining fat was removed from spleens and splenic tissue was mechanically disintegrated using the back of a syringe. Cells were pelleted at 400 g and erythrocytes were lysed using ACK Lysing Buffer (Lonza #10-548E) for 5 minutes at 4°C. Cells were seeded at $10^6$ cells/ml in 48 well plates in a total volume of 500 µl. Cytokines were added as described above for a total of 8 hours.

#### Flow cytometry

Flow cytometry was performed on a BD LSRII flow cytometer. At least 50,000 events were recorded per sample. FCS files were exported by FACSDiva and subsequently gated and compensated in FlowJo (v10.8.0) for single, living and $CD15^+$ (human) and $Ly6G^+$ (murine) cells. Eosinophils were excluded based on high autofluorescence in the live/dead (Pacific Orange) channel **(Supplementary Figure 8)**. Gated events and their median fluorescence intensity values were exported and concatenated into a single cell experiment using CATALYST (v1.16.2; (26)) in R (v. 4.2.0). The dataset was arcsinh transformed using manually determined cofactors (25, 27) and clustered by FlowSOM clustering and Consensus-Plus-Metaclustering. For combined analysis of human and mouse cells, both datasets were mean-centered, scaled and combined into one SingleCellExperiment. Dimensionality reduction was performed using the DiffusionMap algorithm as implemented in the CATALYST package with standard settings. For visualization, a random subset of 1,500 cells per sample were plotted using ggplot2 (v3.3.5; (28)). Principal components were calculated based on the median fluorescence values of the respective marker proteins per sample and plotted using matplotlib (v3.5.1) in python (v3.9.1).

#### Gene Ontology enrichment analysis

We used EnrichR (29-31) to assess enriched gene sets in a given list of genes. The analysis was restricted to terms annotated in GO_Biological_Process_2021

#### Linear Mixed-Effect Model

We used lme4 (v1.1-29; (15)) to fit a linear mixed-effects model (LMM) to $\log_2$(TPM+1) data (filtered as described in **Methods**) in order to validate the core inflammation program derived from Fisher's combined test. The linear formulae were defined as $full = condition + 1|study$ and $reduced = 1|study$, where the variable to test for was *condition*, and the *study* was used as the covariate that was considered to be the random effect. Subsequently, it was tested for each gene in the expression matrix. During the fit, the estimates were chosen to optimize the log-likelihood. We retrieved β as an estimate for the $\log_2$(FC) from the full model and subsequently performed a likelihood ratio test to compare the *full* with the *reduced* model and to retrieve the respective *P*-values. *P*-values were then adjusted using the Benjamini-Hochberg procedure.

#### $\pi_1$-statistic

Using the qvalue-package (v2.28.0; (9, 32, 33)), we calculated the $\pi_1$-statistic (1- $\pi_0$) as an estimated proportion of truly significantly differentially expressed genes for a given set of *P*-values. To account for a selection of genes potentially biased towards low p values when testing for the replicability of DEGs between studies, qvalue-calculation was implemented via the qvalue_truncp function.

#### Gene expression modules using WGCNA

For WGCNA (33) analysis, we selected the same samples, that were used for differential expression testing. We accounted for batch effects by correcting gene counts using ComBat-Seq (sva v3.44.0; (34)) . From batch corrected counts, we calculated TMM-normalized log2 counts per million that were then used as input for WGCNA. The network was constructed as signed network, using a soft thresholding power of 8, minimum module size of 30 and a merge cut height of 0.25. Modules with more than 1000 genes were removed from subsequent analyses.

#### Statistical analyses

Correlations indicated on scatter plots represent Pearson's *R* (Pearson's correlation coefficient) with their respective *P*-value. For comparisons of the mean, we used the Mann-Whitney *U* test (two groups) or Kruskal-Wallis *H* test (three groups), if the Shapiro-Wilk test indicated non-normality in at least one group. When multiple comparisons were performed, *P*-values and/or asterisks indicate adjusted *P*-values using the Holm-Bonferroni method, unless stated otherwise. For pairwise comparisons, the Mann-Whitney *U* test was used post-hoc, taking multiple comparisons into account using the Holm-Bonferroni method. To test for categorical associations, we used Fisher's exact test. Asterisks represent the following *P*-value ranges: $P > 0.05$, ns. $0.01 < P \leq 0.05$, *. $0.001 < P \leq 0.01$, **. $0.0001 < P \leq 0.001$, ***. $P \leq 0.0001$, ****.

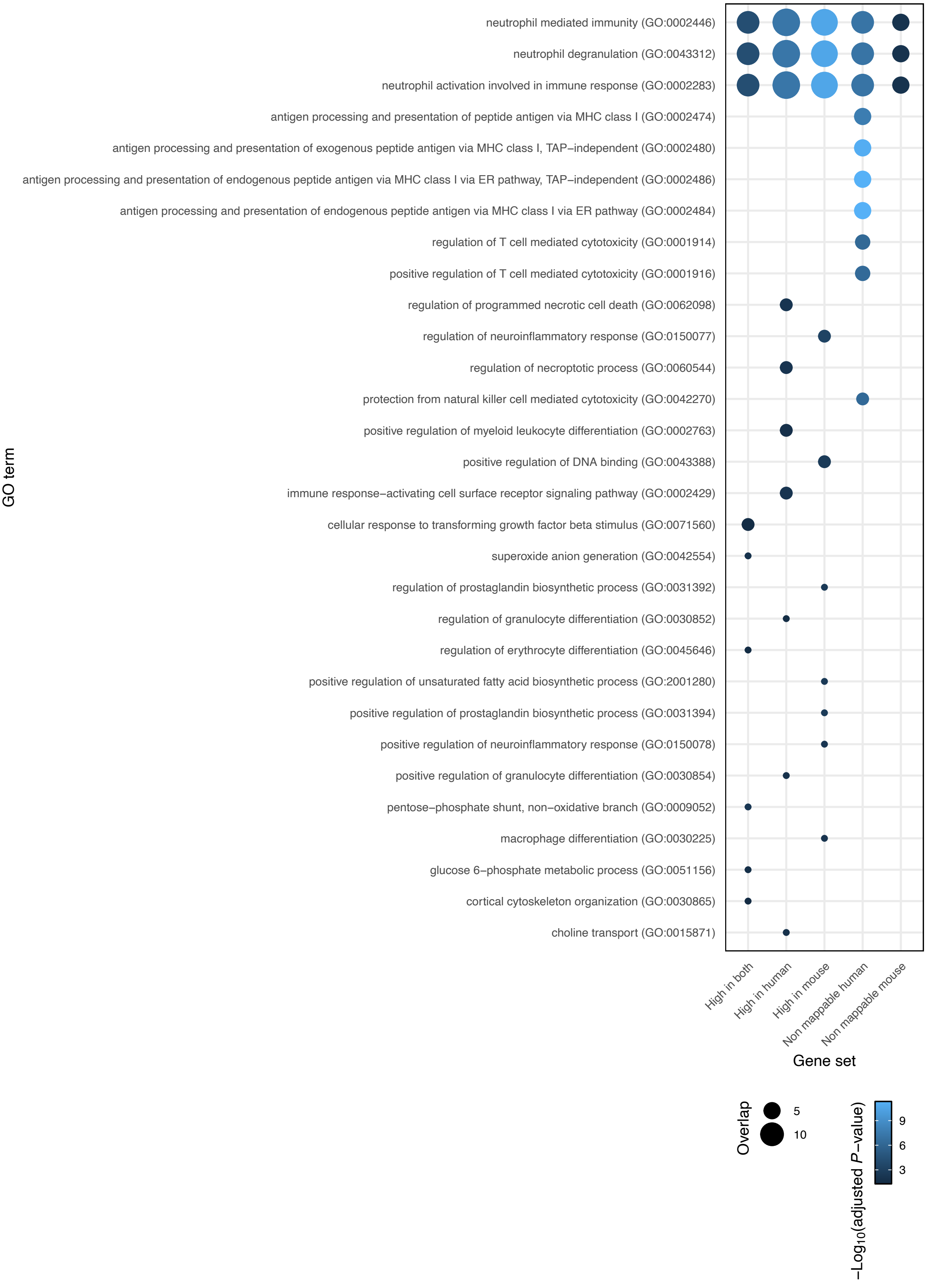


**Supplementary Figure 1. Enrichment analysis for gene clusters in homeostasis.**
Gene Ontology enrichment analysis results for five clusters showing different expression profiles between human and murine neutrophils. Analysis was restricted to the "Biological Process" domain as defined by the Gene Ontology Consortium (35, 36). Only significant enrichments are plotted as dots. Dot size corresponds to overlapping gene set size, color to the negative logarithm of adjusted P-values. GO terms are ranked according to their mean overlap across all significant hits from top to bottom.

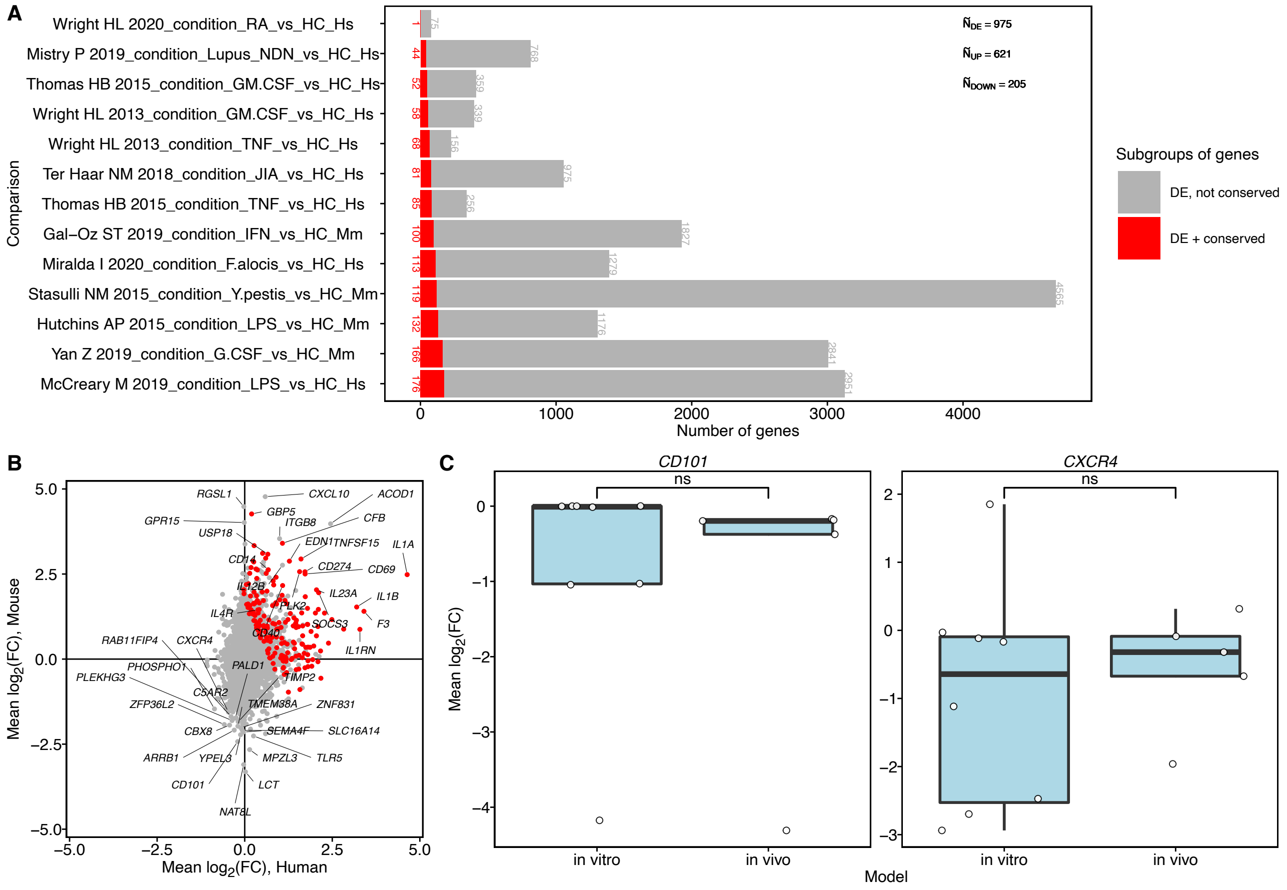


**Supplementary Figure 2. Transcriptional diversity in inflamed neutrophils.**
**(A)** Bar chart indicating the number of genes differentially expressed in each comparison, and the subset of genes that are differentially expressed and part of the core inflammation program. Top right, median number of genes differentially expressed, and the subset of upregulated and downregulated genes. **(B)** Scatter plot of mean $log_2$ fold changes in human versus in mouse. Red, core inflammation program genes. We labeled the genes with the 20 highest and 20 lowest mean expression values as well as genes that were selected for validation (Figures 6 and 7). **(C)** Individual log2 fold changes for CD101 and CXCR4. Here, in vitro and in vivo models of inflammation were separated.

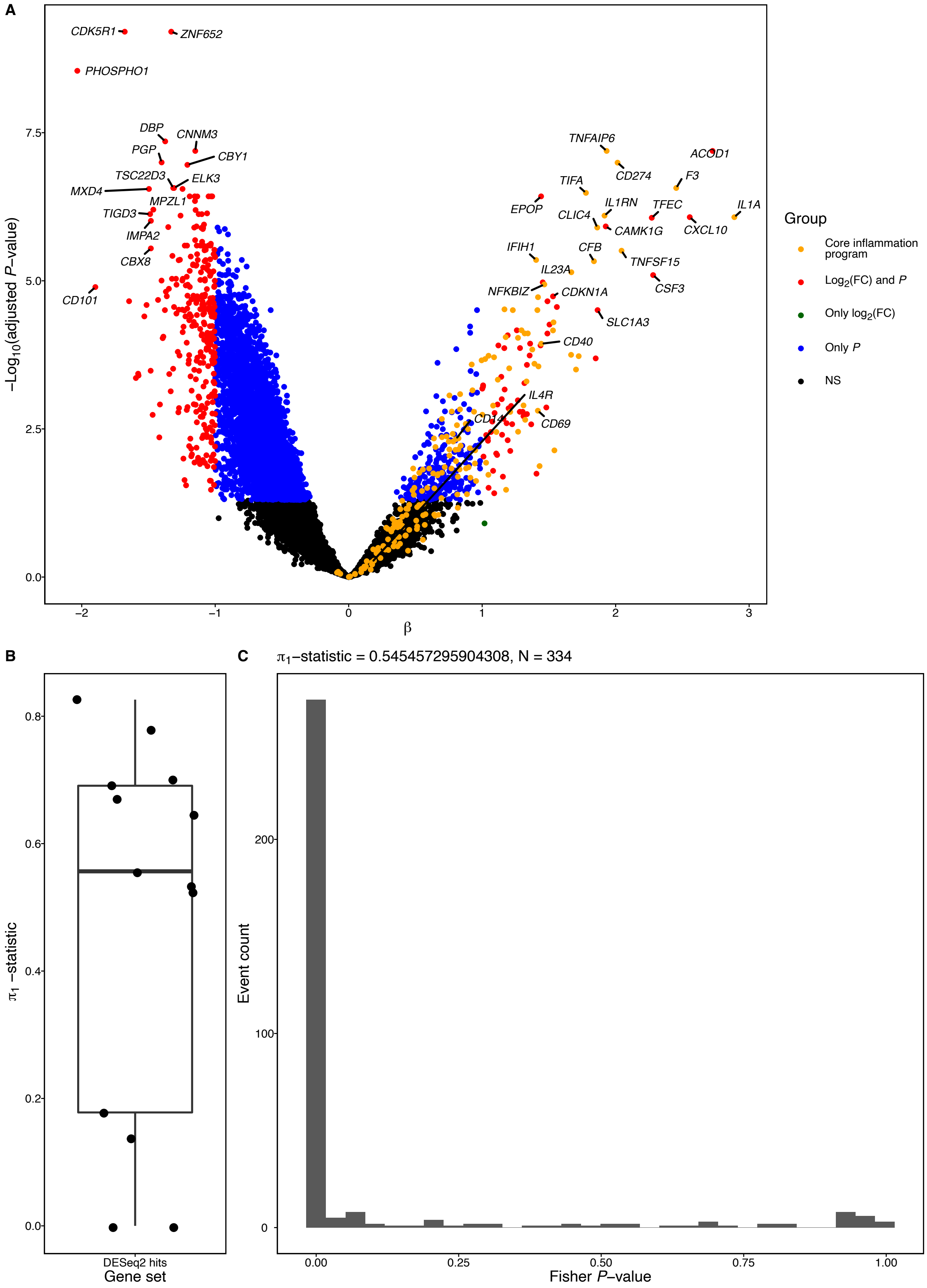


**Supplementary Figure 3. High concordance of complementary linear mixed modeling approach with individual differential expression tests and fisher P-values.**
**(A)** Volcano plot showing the differentially expressed genes according to a linear mixed model testing between healthy control and non-healthy samples. N = 109 upregulated, N = 299 downregulated genes in inflamed neutrophils. Core inflammation genes identified by Fisher's combined approach are highlighted in orange. N = 49 genes identified core inflammation genes pass the threshold in our linear mixed model and are shared. **(B)** Boxplot showing the $\pi_1$-statistic of top differentially expressed genes as identified according to the linear mixed model for each individual differential expression test. **(C)** Histogram showing an enrichment of low fisher *P*-values ($\pi_1 \approx 0.55$) for genes that are up- and downregulated according to the linear mixed model.

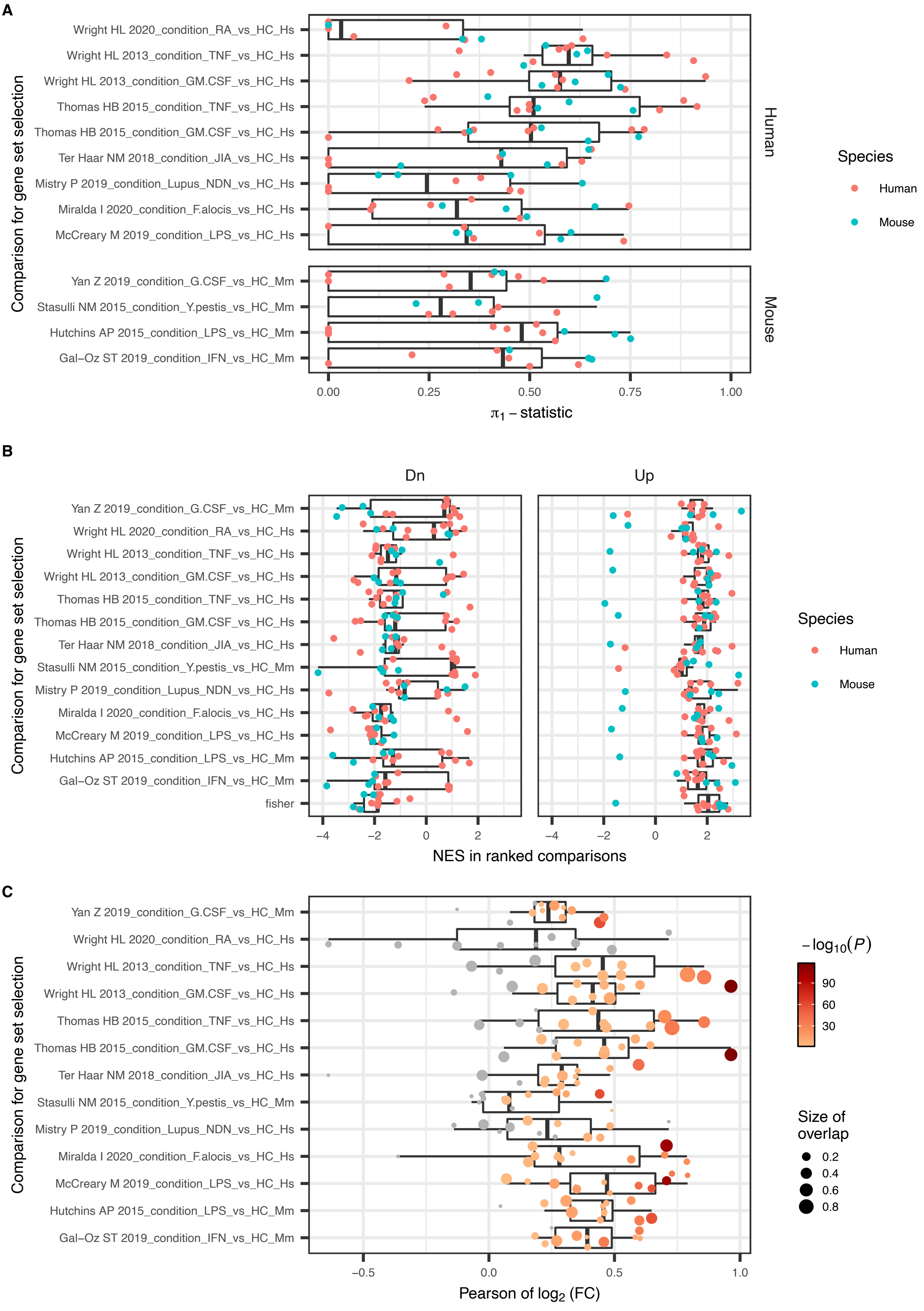


**Supplementary Figure 4. Comparison of differential expression testing between individual studies.**
**(A)** Boxplots depicting the $\pi_1$-statistic of all differentially expressed gene sets (Y-axis) in all other comparisons respectively. Points are colored by species of the respective studies. **(B)** Enrichment scores of differentially expressed gene sets in each comparison (Y-axis) in the ranked differential expression lists of all other comparisons respectively. Points are colored by species of the respective comparison. **(C)** Pearson's *r* of $\log_2$ fold changes of differentially expressed genes in each comparison that are also differentially expressed in other comparisons. Point size corresponds to overlap of differentially expressed genes, color to the negative logarithm of the P-value obtained from a Pearson product-moment correlation test. 6 correlations could not be determined due to a low overlap in differentially expressed genes.

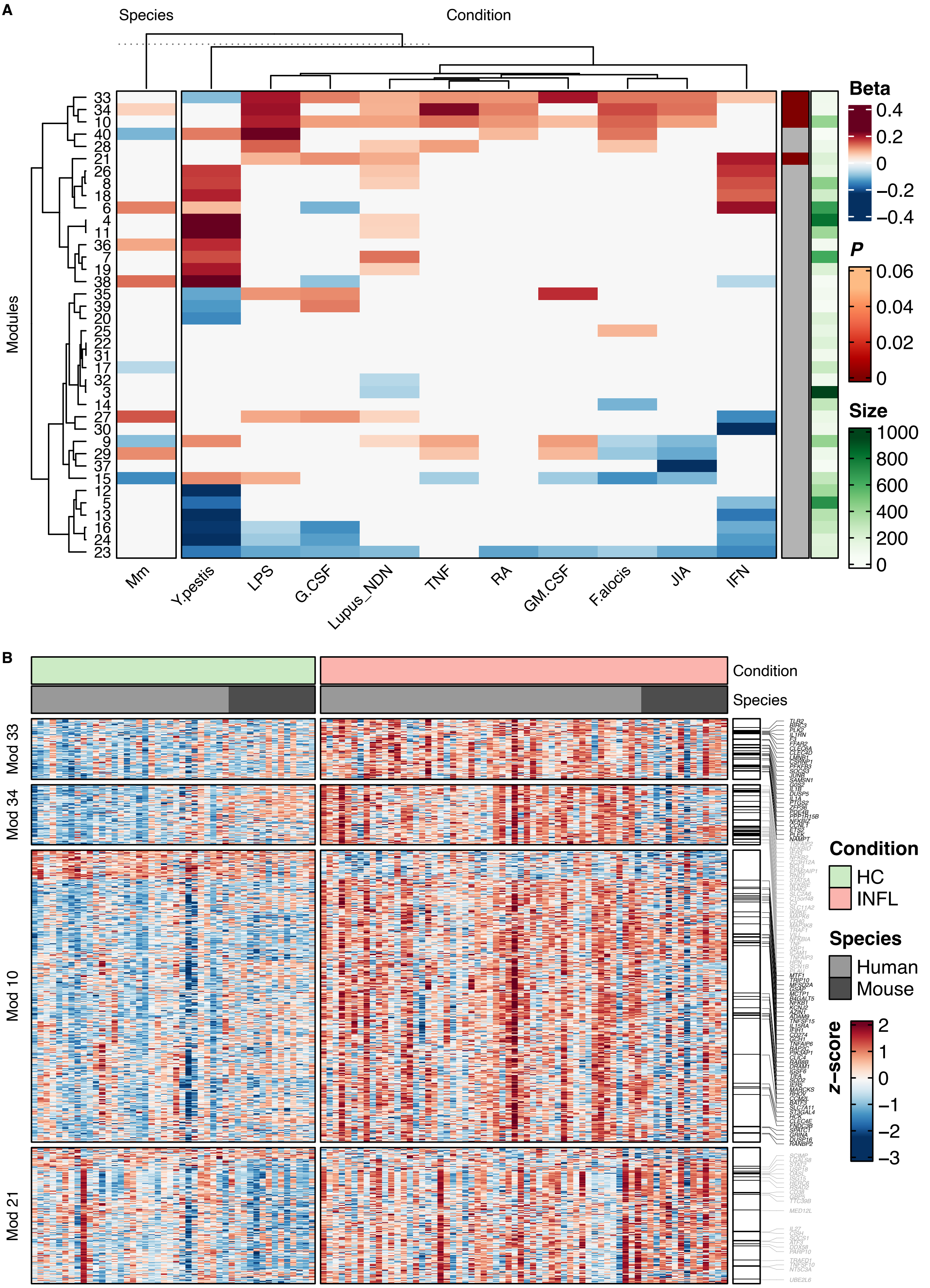


**Supplementary Figure 5. WGCNA analysis across all studies used for differential expression analysis.**
**(A)** Association of gene modules with each experimental condition and species. Eigengenes, a proxy of module expression, for each module were modeled as a combination of condition and species, using a linear model. The heatmap shows effect size estimates resulting from this modeling procedure. Effect size estimates resulting from non-significant associations were set to 0 to allow hierarchical clustering. For each module, a Fisher's exact test was performed to test for association with the core inflammatory response signature. Adjusted *P* values are annotated on the right, gray indicates non-significance. Additionally, module size, defined as the number of genes assigned to that module, is annotated. **(B)** Relative expression profiles (z-score) of genes in modules that are significantly associated with the fisher core inflammatory response signature. Rows are genes, columns are samples. Relative expression was calculated after batch correction for each study (ComBat). Black bars on the right of the expression profiles indicate gene membership in the fisher core signature.

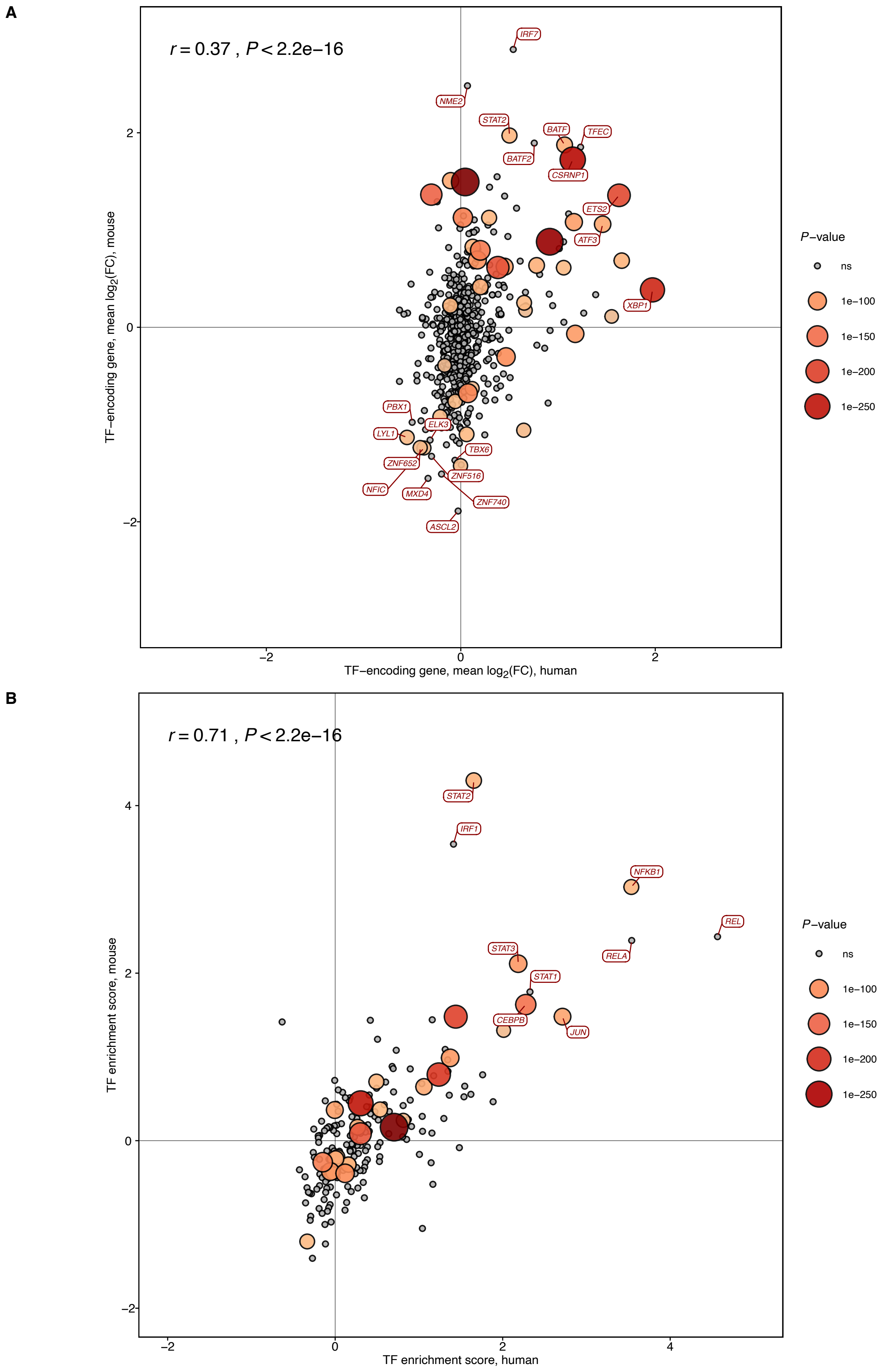


**Supplementary Figure 6. Species-specific Transcription Factor Enrichment Analysis.**
**(A)** Scatter plot of mean $log_2$ fold changes per species for TF encoding genes. The collections of TF-encoding genes was retrieved from DoRothEA (v1.8.0; (22)). Genes with the 10 highest and 10 lowest sums of expression were labeled. **(B)** Scatter plot of transcription factor enrichment scores calculated per species using decoupleR (v2.2.2; (23)). The transcription factors with the 10 highest scores were labeled.

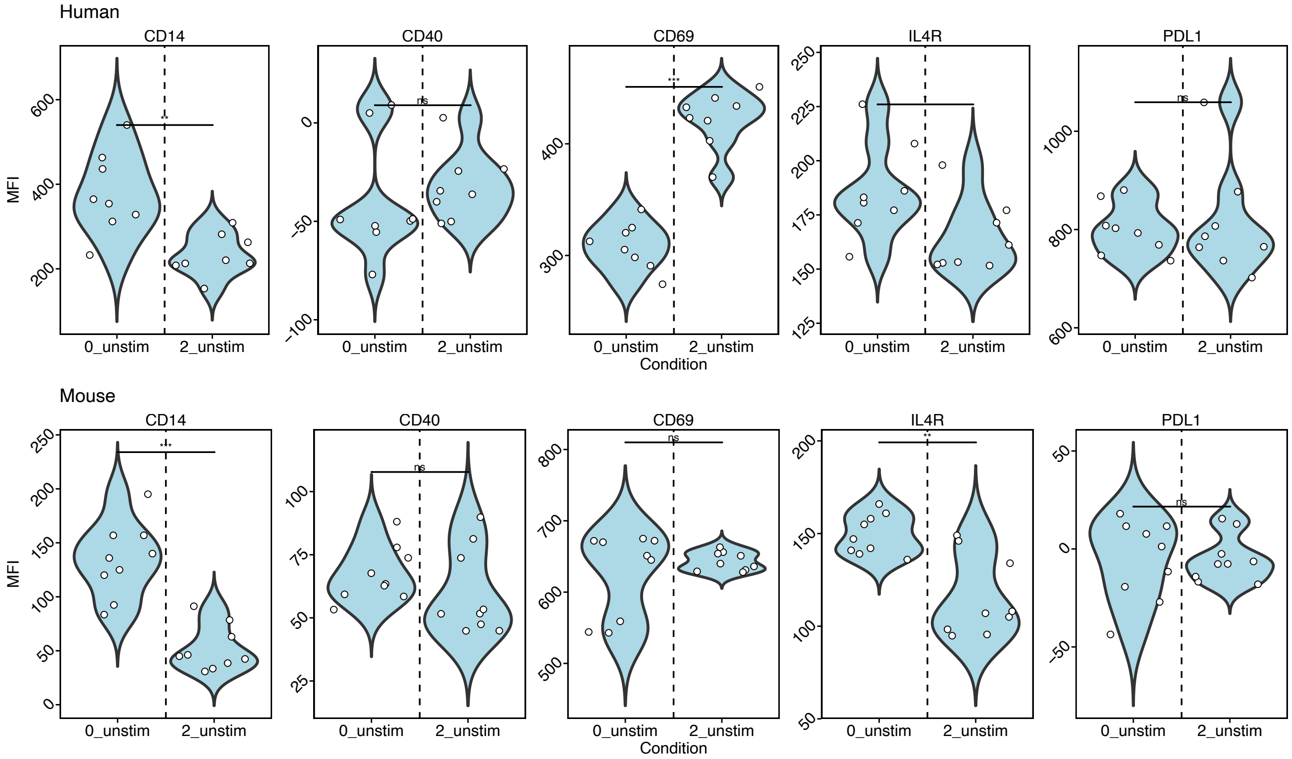


**Supplementary Figure 7. Flow cytometry analysis of neutrophil aging in vitro.**
MFI values of surface proteins depicted in Figure 6C, with and without 2 days of cell culture.

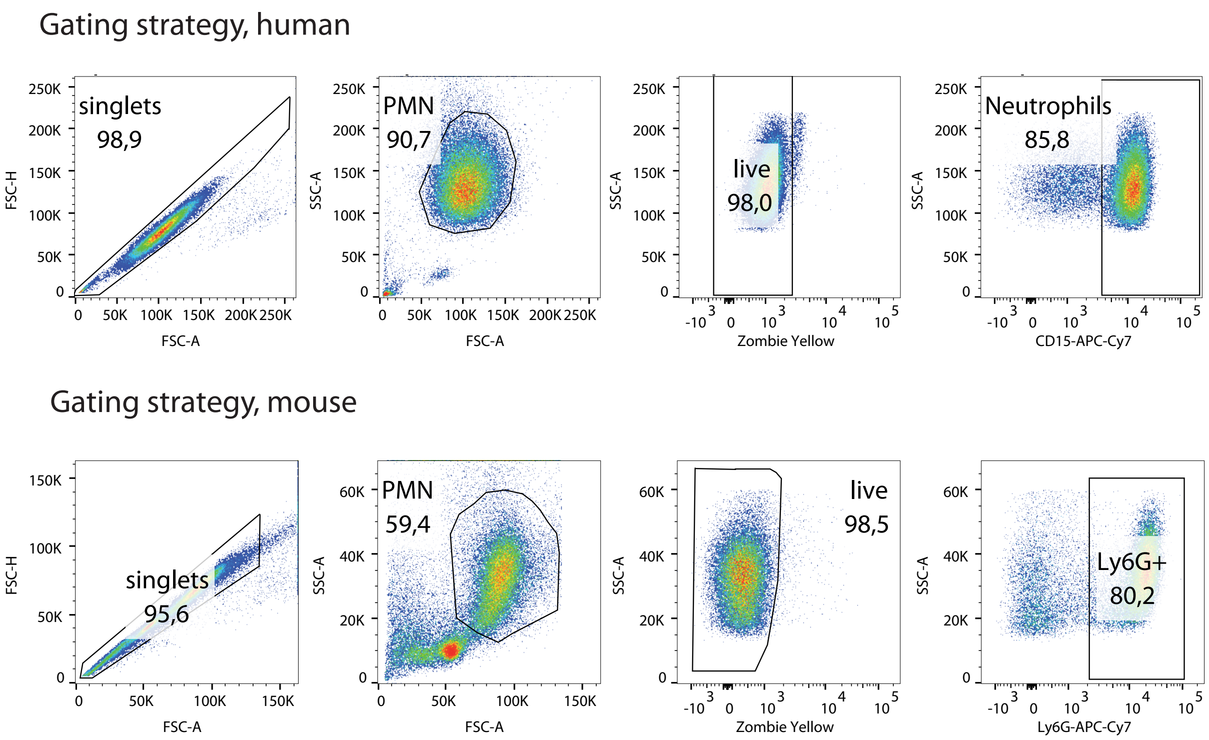


**Supplementary Figure 8. Flow cytometry analysis of murine and human neutrophils.**
Gating strategy for human (top) and murine (bottom) neutrophils.
